# Epigenetic Reactivation of CNS Endothelial Developmental Programs Triggers Adult Brain Angiogenesis, Promotes Post-Stroke Revascularization and Neuronal Regeneration

**DOI:** 10.1101/2025.08.15.670554

**Authors:** Sithara Thomas, Cikesh Chandran, Lalit K. Ahirwar, Arif O Harmanci, Shafeeque M Chathathayil, Nabil J Alkayed, Ari C Dienel, Spiros L. Blackburn, Devin W. McBride, Kumar. T Peeyush

## Abstract

Therapeutic angiogenesis is essential for regenerating brain tissue damaged by stroke, yet it remains an unmet clinical challenge. During brain development, pro-angiogenic genes drive the formation of vascular networks, with their expression tightly regulated in later stages. We found that in adult CNS endothelial cells (ECs), angiogenesis-related genes are epigenetically silenced through histone deacetylase 2 (HDAC2) and the polycomb repressive complex 2 (PRC2). Conditional deletion of *Hdac2* in ECs reactivated pro-angiogenic signaling, including Wnt/β-catenin target genes, leading to functional neovascularization with preserved blood-brain barrier (BBB) integrity in the adult brain. In contrast, *Ezh2* (PRC2 subunit) deletion reduced vessel density and compromised BBB function. Deletion of *Hdac2* and *Ezh2* immediately after transient ischemic stroke conferred vascular protection by modulating stroke-induced transcriptional programs in CNS ECs. In contrast, delayed deletion, initiated seven days post-stroke, after significant neuronal loss in the infarct region, induced robust revascularization and promoted post-stroke neurogenesis, with differentiation into both excitatory and inhibitory neurons. These findings highlight CNS EC HDAC2 as a promising therapeutic target for inducing adult brain angiogenesis, facilitating revascularization, and supporting neuronal regeneration following stroke.

## Introduction

Therapeutic angiogenesis in the adult brain holds promise for mitigating ischemic or neurodegenerative injury. While VEGF administration can stimulate vascular growth, resulting neovessels are structurally immature, lack functional BBB properties, and exhibit context-dependent efficacy^1–3^. For instance, intraventricular VEGF delivery, 48 hours post-ischemia reduces BBB leakage, whereas early post-stroke administration exacerbates vascular dysfunction^3^. Despite robust angiogenic responses in preclinical models, clinical trials employing VEGF or fibroblast growth factor (FGF) proteins or gene therapies have not translated into meaningful functional or clinical benefits^4,5^. Thus, there is an urgent need for alternative approaches to generate stable, barrier-competent vasculature.

In mice, cortical angiogenesis initiates at embryonic day 9.5 (E9.5), with vessel density rising sharply between E13.5 and E15.5, a period coinciding with blood-brain barrier (BBB) maturation^6,7^. Our prior work revealed that transcriptional silencing of angiogenesis and BBB-associated genes, orchestrated by epigenetic regulators *Hdac2* and Polycomb Repressive Complex 2 (PRC2), is critical for stabilizing the developing vascular network^8^. Notably, these repressive mechanisms persist in adult CNS endothelial cells (ECs), where they suppress pro-angiogenic pathways, including Wnt/β-catenin signaling, to maintain cerebrovascular quiescence.

Here, we test the hypothesis that targeted deletion of *Hdac2* and PRC2 (via *Ezh2*) in adult CNS ECs reverses epigenetic repression of developmental angiogenic programs, enabling functional neovascularization. Further, we evaluate whether such vessels enhance neuroprotection in ischemic stroke. While adult neurogenesis occurs in niches such as the hippocampal subgranular zone (SGZ) and subventricular zone (SVZ), post-stroke neuroblast migration to infarcted regions often culminates in apoptosis due to a hostile microenvironment lacking vascular support^9,10^. We hypothesize that *Hdac2* ablation reprograms the neurovascular niche, coupling angiogenesis with neuronal survival and integration, leading to neuronal regeneration in stroke-damaged brains.

By integrating vascular quantification, transcriptional profiling, and stroke recovery assays, we demonstrate that *Hdac2* deletion (1) reactivates Wnt/β-catenin-driven angiogenesis, yielding BBB-intact neovessels; (2) preserves cerebrovascular integrity post-ischemia; and (3) supports regeneration of excitatory and inhibitory neurons in infarcted tissue. Our findings challenge the paradigm that adult brain angiogenesis inevitably produces dysfunctional vasculature, offering a mechanistic blueprint for epigenetic vascular reprogramming in neurological disorders.

## Results

### CNS EC *Hdac2* deletion promotes angiogenesis and the formation of new functional vessels in the adult brain, while *Ezh2* deletion leads to vascular regression

We previously established that during embryonic development, HDAC2 and PRC2 epigenetically repress critical endothelial genes governing angiogenesis and BBB formation^8^. To investigate whether *Hdac2* and *Ezh2* deletion can re-activate angiogenic programs in the adult brain, we administered five doses of tamoxifen (TM) to 2-month-old *Hdac2^flox/flox^*:*Cdh5CreERT2* or *Ezh2^flox/flox^*:*Cdh5CreERT2* mice (Fig. 1a). Brain tissues were harvested two months after the final TM injection for analysis.

**Figure 1:**
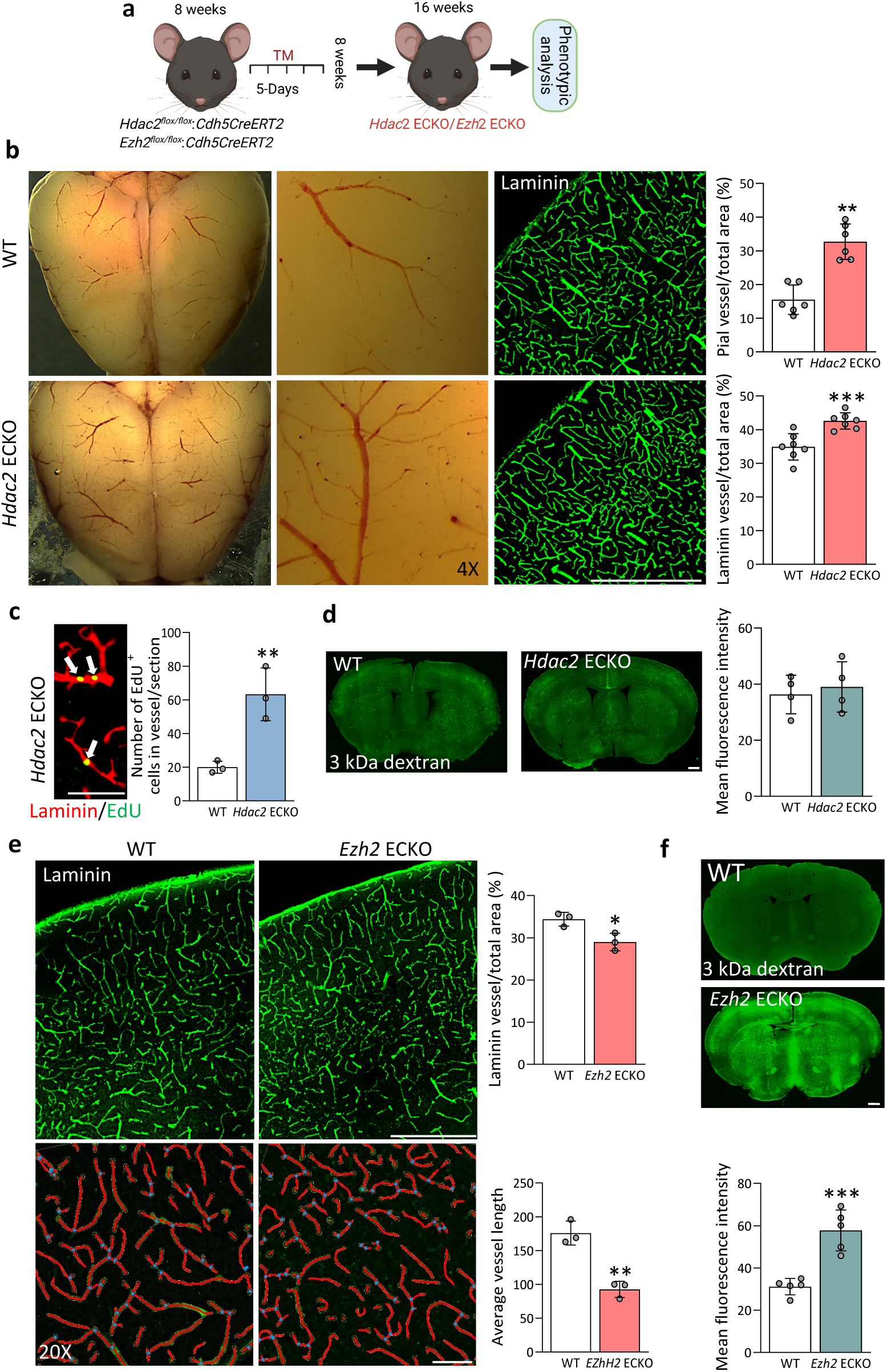
Deletion of endothelial cell *Hdac2* induces neovascularization and *Ezh2* results in vessel regression in adult brain. **(a)** Schematic representation of the experimental plan. Tamoxifen (TM)(75 mg/kg body weight) was administered to *Hdac2^flox/flox^*: C*dh5CreERT2* or *Ezh*2*^flox/flox^*: *Cdh5*^CreERT2^ 8 weeks old mice for generating *Hdac2* ECKO and *Ezh2* ECKO respectively and brains were collected for phenotypic analysis after 8 weeks of final TM injection. **(b)** Representative phase microscopy images of the dorsal surface *Hdca*2 ECKO mice showing a significant increase (**p ≤0 .01 vs. WT. N=3, one hemisphere from each mouse brain was selected for quantification.) in pial vessel density and cortical vessels stained with laminin showing significantly increased vascular density in *Hdac2* ECKO compared to the WT (***p ≤ 0.001 vs. WT. N=6/group and *Hdac2* ECKO N=7/group). Scale bar: 500µm. **(c)** Staining for vessels using laminin and proliferative marker EdU showing proliferating cells in *Hdac2* ECKO mice cortical vessels as indicated by white arrows. Scale bar 500µm. Quantification of EdU^+^ cells in vessels showed significantly increased proliferating cells in *Hdac2* ECKO mice compared to WT (**p ≤ 0.01 vs WT. N=3/group: quantifications were performed on matching sections from independent brains). **(d)** BBB permeability assay using 3kDa-FITC dextran showed no leakage of tracer into brain parenchyma in *Hdac2* ECKO mice (N=4/group: quantifications were performed on matching sections from independent brains ). Scale bar: 500µm. **(e)** Laminin staining displayed a reduction in cortical vessel density in *Ezh2* ECKO mice compared to WT (*p ≤ 0.05 vs WT. N=3/group: quantifications were performed on matching sections from independent brains). Scale bar 500µm. The vessel tracing in the 20X image using AngioTool showed a significantly reduced vessel length in the *Ezh2* ECKO cortex. (**p ≤ 0.01 vs WT. N=3/group quantifications were performed on matching areas of the cortex from independent brains). Scale bar: 5µm**. (f)** BBB permeability assay using 3kDa tracer shows significant leakage into the brain parenchyma of *Ezh2* ECKO mice (***p ≤ 0.001 vs WT. N=5/group: quantifications were performed on matching sections from independent brains). Scale bar: 500 µm. 10x images are acquired and merged using tile scanning. All data are presented as mean ± SD.

We found that *Hdac2* deletion in CNS ECs induces robust angiogenesis in both pial and parenchymal vessels. To assess pial vessel density, non-perfused brains were imaged immediately post-dissection under a light microscope, revealing a marked increase in pial vasculature density in *Hdac2* ECKO mice compared to wild-type (WT) controls, with no signs of hemorrhage (Fig. 1b). Immunostaining with laminin and isolectin B4 (IB4) on 35 µm brain sections further confirmed a significant increase in cortical capillary density (Fig. 1b, Fig. 1—figure supplement 1a), with a similar elevated vessel density observed in the hippocampus, indicating that angiogenesis is broadly distributed across brain regions (Fig. 1—figure supplement 1b).

To confirm EC proliferation in the CNS, EdU (300 µg/kg) was administered for three consecutive days prior to sacrifice. Quantification of EdU⁺ vascular cells revealed a significant increase in *Hdac2* ECKO brains compared to WT controls (Fig. 1c). A key concern in neovascularization is the potential compromise of BBB integrity. To assess this, 3 kDa Alexa Fluor 488-conjugated dextran was perfused, and fluorescence intensity in the brain parenchyma showed no significant difference between *Hdac2* ECKO and WT, indicating intact BBB function in newly formed vessels (Fig. 1d). To further evaluate vascular integrity, we analyzed tight junction (TJ) proteins and associated cell types. ZO-1 staining revealed a significant increase in TJ protein expression in *Hdac2* ECKO vessels (Fig. 1—figure supplement 1c), while CLDN5 was robustly expressed in EdU⁺ vessels, confirming junctional maturation in proliferating endothelium. Pericyte coverage, assessed by PDGFRβ staining, appeared comparable between *Hdac2* ECKO and WT vessels (Fig. 1—figure supplement 1e). Astrocyte analysis via ALDH1 and GFAP revealed a marked increase in astrocyte density in *Hdac2* ECKO brains, accompanied by elevated AQP4 staining, indicating enhanced astrocyte–vascular interactions (Fig. 1—figure supplement 1f). Finally, laminin staining of peripheral organs (liver, lung, and kidney) showed no detectable differences in vascular structure or tissue morphology between genotypes, suggesting CNS-specific angiogenic activation (Fig. 1—figure supplement 1g).

To evaluate the functional impact of *Hdac2* deletion, we conducted behavioral assays, open field, Y-maze, and Barnes maze, to assess locomotor activity, spatial memory, and cognitive performance^11–13^ in *Hdac2* ECKO mice eight weeks post-TM injection. *Hdac2* ECKO mice showed normal behavior, indicating that *Hdac2* deletion-driven angiogenesis does not impair baseline locomotor or cognitive functions (Fig. 1—figure supplement 1h).

In contrast, *Ezh2* deletion resulted in a significant reduction in vascular density and vessel length in the brain two months post-deletion, as compared to WT controls (Fig. 1e). BBB integrity was compromised in Ezh2 mutants, as evidenced by extensive leakage of 3 kDa dextran (Fig. 1f). Behaviorally, *Ezh2* mutants demonstrated hyperactivity, with increased cumulative duration in the center zone and greater total distance traveled in the open field test. However, no significant deficits were observed in Y-maze or Barnes maze performance relative to WT mice (Fig. 1—figure supplement 2).

### Deletion of *Hdac2* and *Ezh2* activates angiogenesis and BBB genes in CNS ECs, with *Hdac2* deletion inducing a broader gene activation profile than *Ezh2*

To elucidate the transcriptomic basis underlying the different phenotypes of *Hdac2* ECKO and *Ezh2* ECKO mice, we performed mRNA sequencing of cortical ECs sorted two months post-TM injection (Fig. 2a). Live CD31^+^, CD41^−^, CD45^−^ cortical ECs were isolated, and *Hdac2*^flox/flox^ mice treated with TM served as controls. Differential expression analysis identified 2,254 upregulated and 2,297 downregulated genes in *Hdac2* ECKO ECs (p ≤ 0.05; Fig. 2b). Gene ontology analysis revealed significant enrichment in vascular categories, including angiogenesis, sprouting angiogenesis, EC proliferation, migration, and Wnt signaling (Fig. 2c). *Hdac2* deletion broadly activated genes in all categories, with some repression observed across all except EC migration. BBB-associated gene sets, cell-cell junction, transporters, and barrier maintenance, also displayed substantial transcriptional remodeling, with both up- and downregulated genes (Fig. 2d).

**Figure 2:**
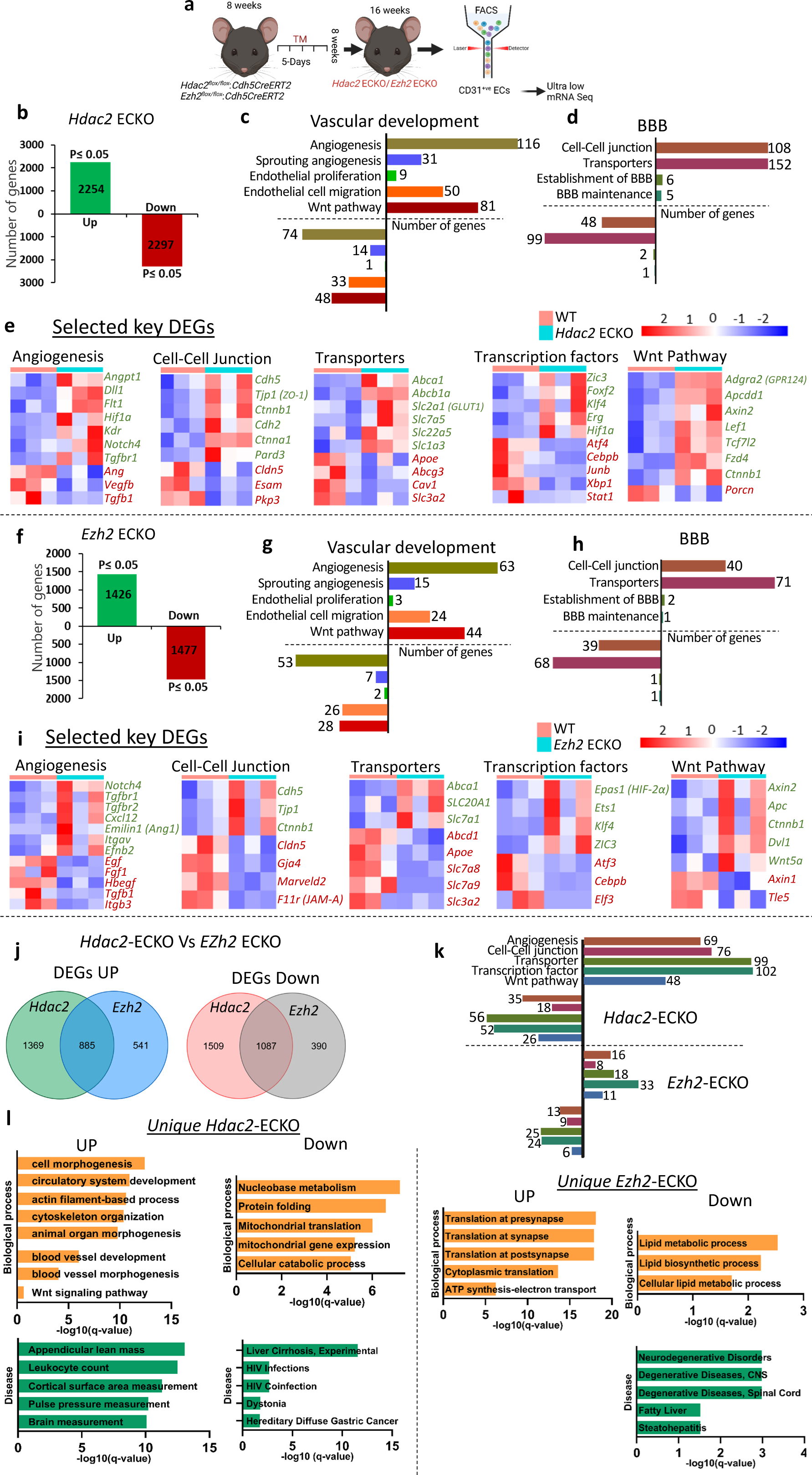
Gene expression profile of CNS ECs after Hdac2 and *Ezh2* deletion. (**a**) Schematic representation of the experimental design. Tamoxifen (75 mg/kg body weight) was administered to *Hdac2^flox/flox^*:*Cdh5CreERT2* or *Ezh2^flox/flox^*:*Cdh5CreERT2* mice at 8 weeks of age. After 8 weeks, live CD31+/CD41-/CD45-brain ECs were isolated by FACS sorting and subjected to ultra-low mRNA sequencing. (**b, f**) Differential gene expression analysis showing the number of significantly upregulated and downregulated genes (p ≤ 0.05 N=3/group) in cortical ECs of *Hdac2* ECKO (b) and *Ezh2* ECKO (f) mice compared to WT controls. (**c, g**) Classification of differentially expressed genes (DEGs) into vascular development-related gene ontology (GO) terms associated with the observed phenotypes in *Hdac2* ECKO (c) and *Ezh2* ECKO (g) mice. (**d, h**) Subclassification of DEGs based on BBB-related functional categories in *Hdac2* ECKO (d) and *Ezh2* ECKO (h) mice. (**e, i**) Heatmap showing selected key DEGs categorized into genes related to angiogenesis, cell-cell junctions, transporters, transcription factors, and the Wnt signaling pathway in *Hdac2* ECKO (e) and *Ezh2* ECKO (i) mice. (**j**) Comparative analysis of the number of DEGs (upregulated and downregulated) that are unique or shared between *Hdac2* ECKO and *Ezh2* ECKO ECs. (**k**) Classification of unique DEGs in *Hdac2* ECKO and *Ezh2* ECKO ECs based on GO terms related to angiogenesis, BBB function, and Wnt signaling. (**l**) Top 5 biological processes and diseases associated with the unique upregulated and downregulated genes in *Hdac2* ECKO and *Ezh2* ECKO mice. For *Hdac2* ECKO, additional relevant terms outside the top 5 are also displayed. No significant disease associations were found among the uniquely upregulated genes in *Ezh2* ECKO mice.

*Hdac2* deletion upregulated angiogenic regulators (*Angpt1*, *Dll1*, *Flt1, Kdr (Vegfr2)*), TJ genes (*Cdh5*, *Tjp*1), transporters (*Abca1*, *Slc2a1*), BBB-related transcription factors (*Zic3*, *Foxf2*), and Wnt components and targets (*Adgra2*, *Apcdd1*, *Axin2*, *Lef1*, *Fzd4*, *Ctnnb1*) (Fig. 2e). Concomitantly, several genes were downregulated, including angiogenesis-related (*Ang*, *Vegfb*), tight junction (*Cldn5*, *Esam*), transporters (*Apoe*, *Cav1*), and transcriptional regulators (*Atf4*, *Cebpb*). Top DEGs and enriched biological processes are shown in Fig. 2 – figure supplement 1a. Upregulated genes were associated with neurogenesis, morphogenesis, and vascular development, while downregulated genes were enriched in protein catabolism, folding, and trafficking pathways. We further validated a significant increase in Vegfr2 expression in *Hdac2* ECKO CNS vessels compared to WT (Fig. 2 – figure supplement 1b).

In *Ezh2* ECKO ECs, 1,426 genes were upregulated and 1,477 downregulated (p ≤ 0.05; Fig. 2f). Similar categorization of DEGs revealed activation and repression across angiogenesis and BBB-related terms (Fig. 2g–h). Upregulated angiogenesis genes included *Notch4*, *Emilin1*, and *Itgav*, while growth factor genes such as *EGF* and *FGF1* were downregulated. Ezh2 deletion also induced *Cdh5* and *Tjp1* expression but suppressed *Cldn5*, *Gja4*, and *Marveld2*. Transporter genes such as *Abca1* and *Slc7a1* were upregulated, while *Abcd1* and *Apoe* were downregulated. Transcription factors *Epas1*, *Klf4*, and *Zic3* were activated, whereas *Atf3* and *Cebpb* were suppressed. Wnt signaling genes, including *Axin2*, *Apc*, and *Ctnnb1*, were upregulated, while *Axin1* and *Tle5* were repressed (Fig. 2i). The top 20 DEGs and associated biological processes are presented in Fig. 2 – figure supplement 1c.

To further refine the molecular basis underlying the distinct vascular phenotypes of *Hdac2* ECKO and *Ezh2* ECKO mice, we directly compared their transcriptomic profiles to identify uniquely regulated gene sets. This revealed that *Hdac2* ECKO uniquely upregulated 1,369 genes and downregulated 1,509 genes, whereas *Ezh2* ECKO uniquely upregulated 541 and downregulated 390 genes relative to *Hdac2* ECKO (Fig. 2j). Classification of these differentially expressed genes (DEGs) based on angiogenesis, BBB, and Wnt pathway terms showed a markedly greater alignment of *Hdac2* ECKO-specific genes with these categories, consistent with its robust pro-angiogenic phenotype (Fig. 2k). The top 20 unique gene set for each condition are shown in Fig. 2 – figure supplement 2a, 2b. Gene ontology enrichment of the *Hdac2* ECKO -specific DEGs identified biological processes such as cell morphogenesis and circulatory system development among the top hits (Fig. 2l). Although not in the top five, terms such as blood vessel development and Wnt signaling were also significantly enriched, supporting the observed vascular remodeling. Downregulated genes were enriched in pathways related to nucleobase metabolism, protein folding, and mitochondrial translation. Disease association analysis revealed that *Hdac2* ECKO-upregulated genes were linked to traits such as appendicular lean mass, leukocyte count, and cortical surface area, while downregulated genes were associated with conditions like liver cirrhosis and HIV infection. In contrast, *Ezh2* ECKO upregulated genes were enriched in synaptic and cytoplasmic translation processes, while downregulated genes were strongly associated with lipid metabolism pathways. Notably, genes downregulated in *Ezh2* ECKO were also enriched for neurodegenerative disease categories, consistent with the altered behavioral phenotype observed, whereas upregulated genes were linked to anemia-related traits (data not shown).

### *Hdac2* and *Ezh2* deletion from ECs prevents stroke-induced brain injury

Given that HDAC2 and PRC2 regulate several critical genes in CNS ECs and their deletion can activate angiogenesis and BBB-related genes, we were eager to explore their effect on stroke-induced vascular injury. To this, we performed transient middle cerebral artery occlusion for 60 minutes, followed by reperfusion for 7 days. To induce *Hdac2* and *Ezh2* deletion, TM was injected one hour after the surgery and continued daily until day 5. Behavioral analysis was performed on day 7, followed by brain harvesting (Fig. 3a).

**Figure 3:**
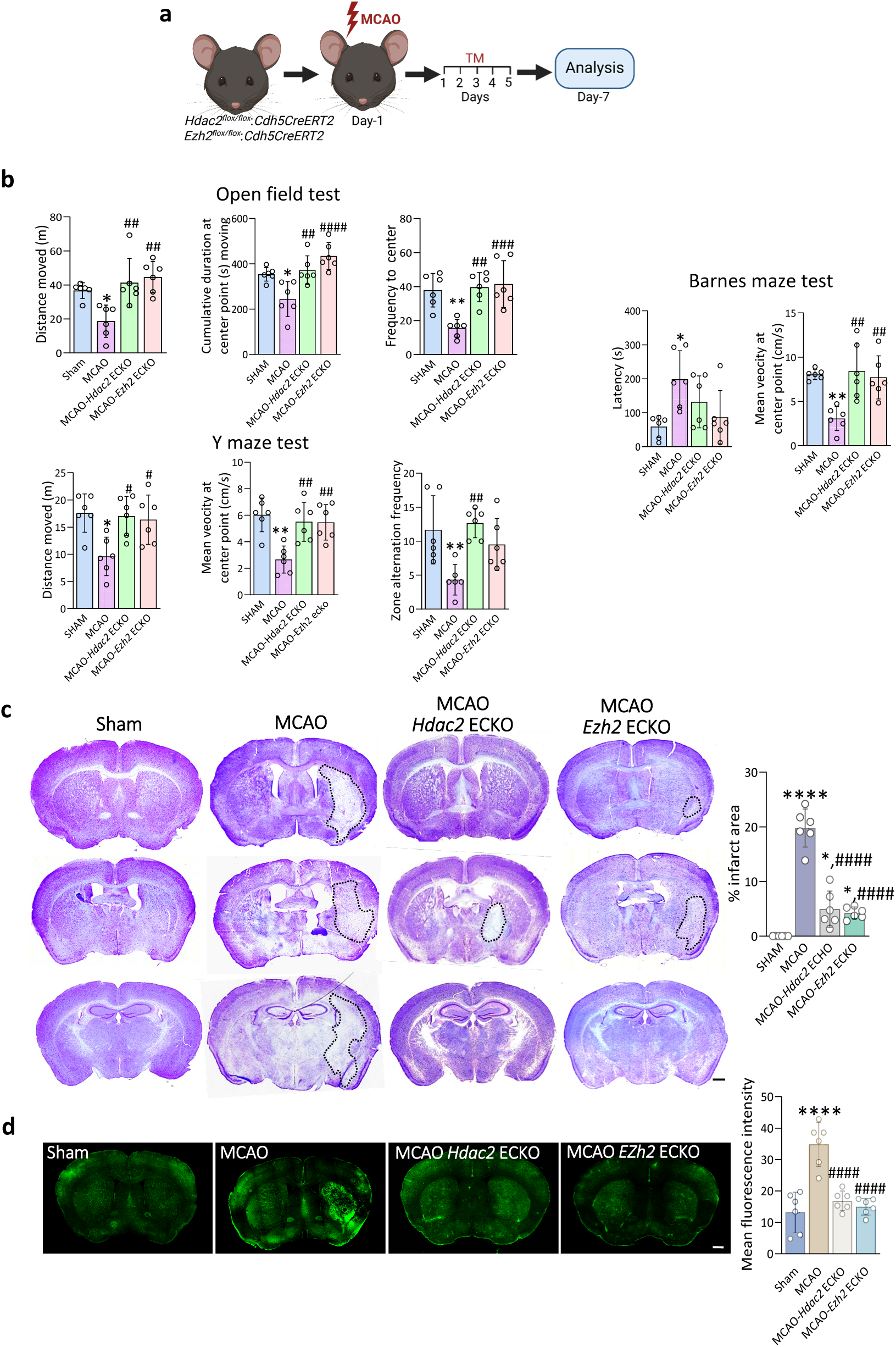
Deletion of *Hdac2* and *Ezh2* from mouse brain endothelial cells improved ischemic stroke outcomes. **(a)** Illustration of experimental strategy. MCAO was induced in 8-weeks-old *Hdac2^flox/flox^*:*Cdhh5CreERT2* and *Ezh2^flox/flox^*:*Cdh5CreERT2* mice and TM was administered one hour after the completion of surgery (75mg/Kg body weight) for five days and mice were sacrificed on 7^th^ day for analysis. **(b)** Behavioral assessments using the Open Field Test (OFT), Barnes Maze Test (BMT), and Y Maze Test (YMT) revealed significant behavioral deficits in MCAO mice, with improvements observed following EC-specific deletion of *Hdac2* or *Ezh2*. In the OFT, MCAO mice displayed reduced total distance traveled, time spent in the center, and frequency of center entries compared to sham controls. Conversely, MCAO-*Hdac2* ECKO and MCAO-*Ezh2* ECKO mice exhibited significant improvements in these parameters, approaching sham levels. In the YMT, MCAO mice showed decreased total distance traveled, mean velocity, and zone alternation frequency relative to controls, while MCAO-*Hdac2* ECKO and MCAO-*Ezh2* ECKO mice demonstrated significant recovery in distance and velocity; MCAO-*Hdac2* ECKO mice additionally showed enhanced zone alternation compared to controls. In the BMT, MCAO mice exhibited increased latency to locate the target hole compared to sham controls, whereas MCAO-*Hdac2* ECKO and MCAO-*Ezh2* ECKO mice showed no significant differences from sham or MCAO groups. Additionally, MCAO mice had reduced mean velocity at the center point, while MCAO-*Hdac2* ECKO and MCAO-*Ezh2* ECKO mice displayed significant increases compared to MCAO alone. (*p ≤ 0.05, **p ≤ 0.01 vs *** p≤ 0.001 vs sham, #p ≤ 0.05, ##p ≤ 0.01, ###p ≤ 0.001 vs MCAO. N=6/group). **(c)** Representative Nissl-stained brain sections from mice subjected to sham surgery, MCAO, MCAO-*Hdac2* ECKO, and MCAO-*Ezh2* ECKO, with infarct areas outlined by a black dotted line. Images were acquired at 10× magnification and reconstructed using tile-scanning microscopy. Scale bar 500µm. Quantification of the infarct area shows a significantly larger infarct in MCAO mice compared to sham, with significantly smaller infarcts in MCAO-Hdac2 ECKO and MCAO-*Ezh2* ECKO mice relative to MCAO. (*p ≤ 0.05, ****p ≤ 0.0001 vs sham, ####p ≤ 0.0001 vs MCAO. N=6/group and matching sections were compared between 6 independent mice brains from each group) **(d)** BBB permeability assay using a 3 kDa FITC-dextran tracer demonstrates leakage into the brain parenchyma in MCAO brain compared to sham. In contrast, MCAO-*Hdac2* ECKO and MCAO-*Ezh2* ECKO mice exhibit no detectable leakage of the 3 kDa FITC-dextran tracer into the brain parenchyma. 10 x images are acquired and merged using tile scanning. Scale bar 500 µm. (****p ≤ 0.0001 vs sham, ####p ≤ 0.0001 vs MCAO N=6/group and N=6, matching sections were compared between 6 independent mice brains from each group). All data are presented as mean ± SD.

To assess locomotor, exploratory, and cognitive deficits post-stroke, we performed three standard behavioral tests 7 days after MCAO, following pre-training. Open field testing showed reduced exploratory activity, slower movement, and increased resting time in MCAO mice. Y-maze testing revealed impaired spatial memory, with decreased distance traveled, fewer arm alternations, and reduced velocity. Barnes maze testing showed increased latency to find the escape hole, indicating spatial learning deficits. *Hdac2* or *Ezh2* ECKO mice exhibited improved performance across all tests compared to MCAO controls (Fig. 3b).

To assess the effect of Hdac2 and Ezh2 deletion on post-stroke brain injury, we performed Nissl staining at day 7 following MCAO. MCAO-only mice showed prominent infarcts consistent with neuronal loss, whereas *Hdac2* ECKO and *Ezh2* ECKO mice had significantly smaller infarct areas, indicating a protective effect (Fig. 3c). Given the association between ischemic stroke and BBB disruption, we assessed BBB integrity using 3 kDa FITC-dextran. MCAO brains exhibited widespread tracer leakage into the parenchyma, confirming BBB breakdown, while *Hdac2* ECKO and *Ezh2* ECKO mice exhibited minimal leakage, confirming preserved BBB function (Fig. 3d).

### *Hdac2* and *Ezh2* deletion mitigates stroke-induced gene expression changes in ipsilateral CNS ECs

MCAO is known to alter vascular gene expression^14–16^. We performed RNA sequencing on CD31⁺CD41⁻CD45⁻ ECs isolated from the ipsilateral and contralateral hemispheres of sham and MCAO brains, as well as from the ipsilateral side of MCAO-*Hdac2* ECKO and MCAO-*Ezh2*ECKO mice on day 7 post-stroke (Fig. 4a) to determine the transcriptomic changes associated with stroke-induced vascular injury and to assess how *Hdac2* or *Ezh2* deletion alters these gene expression programs. WT ipsilateral CNS ECs post-stroke showed 1,479 differentially expressed genes (DEGs) versus sham (965 upregulated, 514 downregulated; padj ≤ 0.05), confirming a strong ischemia-induced transcriptional response (Fig. 4b). In contrast, the contralateral hemisphere showed minimal changes (28 upregulated, 5 downregulated; Fig. 4–figure supplement 4a,b) indicating the response is unilateral. *Hdac2* ECKO and *Ezh2* ECKO mice post-stroke had significantly fewer DEGs, 349 (151 up, 198 down) and 321 (69 up, 252 down), respectively. Comparison with WT MCAO DEGs revealed limited overlap: 83 genes in *Hdac2* ECKO (52 up, 31 down) and 32 in *Ezh2* ECKO (14 up, 18 down) (Fig. 4b). Enrichment analysis of upregulated genes in WT MCAO mice revealed associations with biological processes such as apoptosis, inflammation, cellular stress, and angiogenesis. Conversely, downregulated DEGs were associated with processes like circulatory system development, neuron development, cell adhesion, vascular development, and Wnt signaling (Fig. 4c).

**Figure 4:**
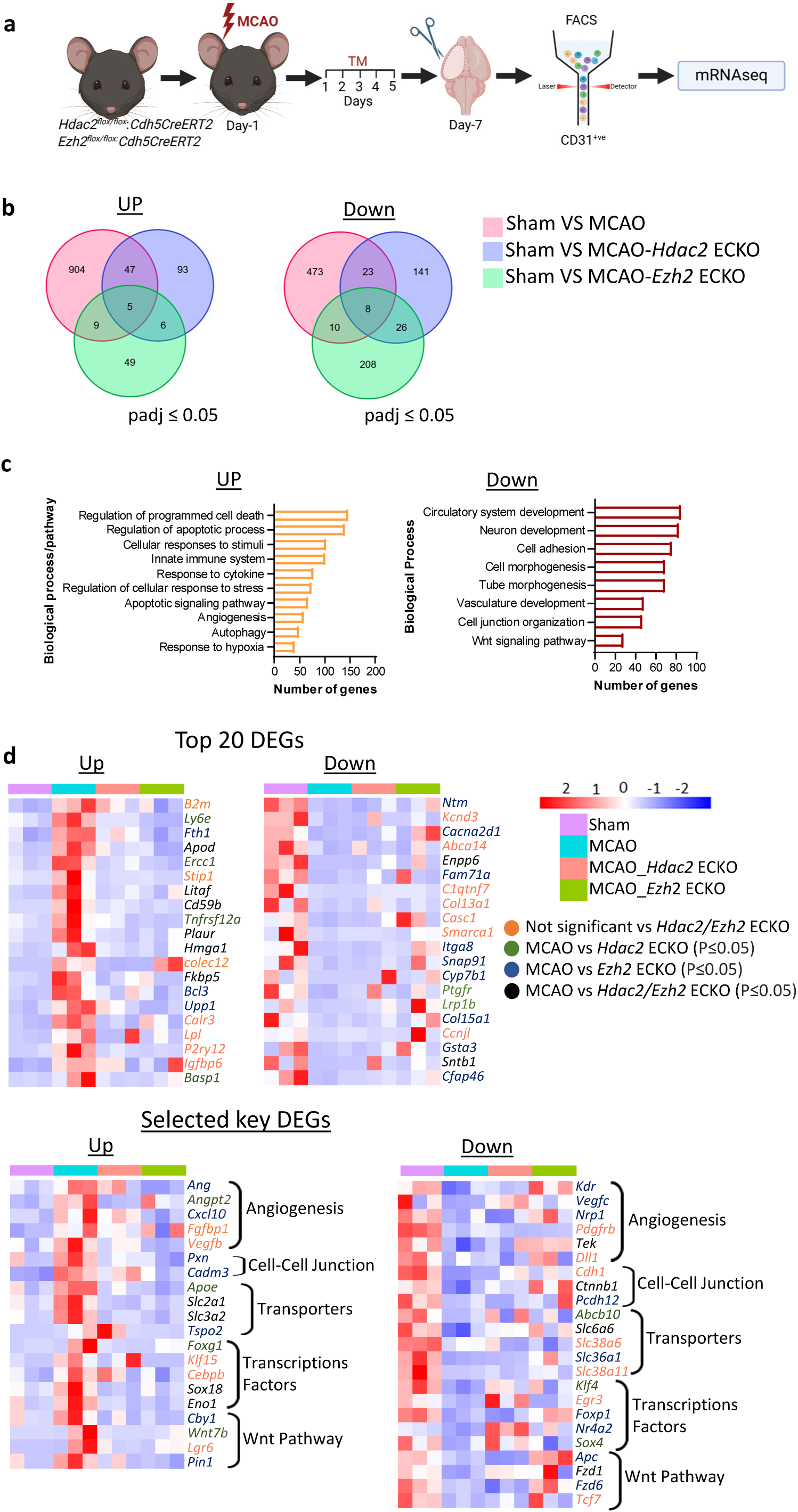
MCAO alters CNS EC gene expression in the ipsilateral hemisphere, while EC-specific deletion of *Hdac2* and *Ezh2* attenuates MCAO-induced gene expression changes. **(a)** Experimental strategy schematic. MCAO was induced in 8-week-old WT, *Hdac2^flox/flox^*:*Cdh5CreERT2*, and *Ezh2^flox/flox^*:*Cdh5CreERT2* mice. Tamoxifen (75 mg/kg) was given one hour post-surgery for 5 days. Mice were sacrificed on day 7 post-MCAO. Ipsilateral cortical ECs were isolated and FACS sorted (CD31^+^/CD41^−^/CD45^−^) for ultra-low-input mRNA sequencing. **(b)** RNA sequencing results showing the number of upregulated and downregulated genes (padj ≤ 0.05 N=3/group) in MCAO, MCAO-*Hdac2* ECKO, and MCAO-*Ezh2* ECKO mice relative to sham mice. **(c)** Enrichment analysis of upregulated and downregulated genes in MCAO mice, highlighting the top 10 associated biological processes. (**d**) Heatmap of the top 20 DEGs in MCAO mice, ranked by fold change (padj ≤ 0.05 N=3/group) relative to sham, with corresponding expression levels in MCAO-Hdac2 ECKO and MCAO-*Ezh2* ECKO mice compared to sham. **(e)** Heatmap of DEGs in MCAO vs sham categorized by selected GO terms (angiogenesis, cell-cell junctions, transporters, transcription factors, Wnt signaling), with corresponding expression levels in MCAO-*Hdac2* ECKO and MCAO-*Ezh2* ECKO. All data are presented as mean ± SD.

To determine whether *Hdac2* or *Ezh2* deletion mitigates MCAO-induced CNS EC gene changes, we examined the top 20 upregulated and downregulated genes in WT MCAO mice. Strikingly, only 7 genes in each group remained unchanged in *Hdac2* ECKO or *Ezh2* ECKO-MCAO mice (Fig. 4d, Fig. 4–figure supplement 4c, d.). GO term analysis of angiogenesis, BBB, junctions, transporters, transcription factors, and Wnt signaling revealed that 15 of 20 upregulated and 16 of 23 downregulated genes were significantly regulated by either deletion. Transcriptomic profiling of contralateral cortical ECs in sham vs MCAO-*Hdac2* ECKO, and MCAO-*Ezh2* ECKO mice revealed robust differential gene expression, confirming efficient recombination and transcriptional reprogramming within this period. These differentially expressed genes were strongly enriched for angiogenesis, BBB, and transcription factor activity, indicating that early epigenetic manipulation rapidly engages key vascular and regulatory pathways in ECs (Fig. 4-figure supplement 4e, f).

### β-catenin GOF fails to induce angiogenesis or confer protection against stroke injury in the adult brain

Supporting our prior findings that Wnt target genes are epigenetically silenced in adult central nervous system (CNS) endothelial cells (ECs), *Hdac2* ECKO mice showed upregulation of Wnt targets such as *Axin2* and *Lef1*. Since some studies indicate Wnt pathway activation in adult CNS ECs via β-catenin gain-of-function (GOF)^10,11^, we test the functional impact of this activation on CNS vessels. To this, we used β-catenin gain-of-function (β-GOF) mice (*Ctnnb1^(ex^*^3^*^)flox/flox^*:*Cdh5CreERT2* ) treated with TM for 5 days to induce stabilized β-catenin. Phenotypically, β-cat-GOF mice showed no significant changes in vascular density or morphology compared to WT. BBB integrity remained intact, with no differences in 3 kDa dextran leakage (Fig. 5b). Behavioral testing via open field, Y-maze, and Barnes maze also revealed no significant cognitive or locomotor differences (Fig. 5 - figure supplement 5a). However, transcriptomic analysis of CD31^+^ CNS ECs revealed 1,280 DEGs (534 upregulated, 746 downregulated), including upregulated Wnt-related genes such as *Flt1*, *Gja1*, *Mfsd2a*, and *Apcdd1* and downregulated genes such as *Edn1, Tgfb1, Marveld2, Abcg3, Cav1*, and *Sox17*. Gene ontology analysis revealed alterations in pathways related to angiogenesis, cell junctions, transport, and Wnt signaling (Fig. 5d). The top 20 DEGs and top biological processes, based on fold change (padj ≤0.05) that are altered in β-cat GOF, are given in Fig. 5 - figure supplement 5b. To compare Wnt activation via β-cat GOF with *Hdac2* ECKO, we analyzed transcriptomic overlaps. *Hdac2* ECKO mice upregulated a broader array of Wnt-associated genes; of the 81 Wnt-related genes upregulated in *Hdac2* ECKO, only 14 overlapped with β-cat GOF, while 67 were uniquely regulated (Fig. 5e), including downstream target genes such as *Axin2* and *Lef1*, and upstream components like *Fzd4* and *Gpr124*.

**Figure 5:**
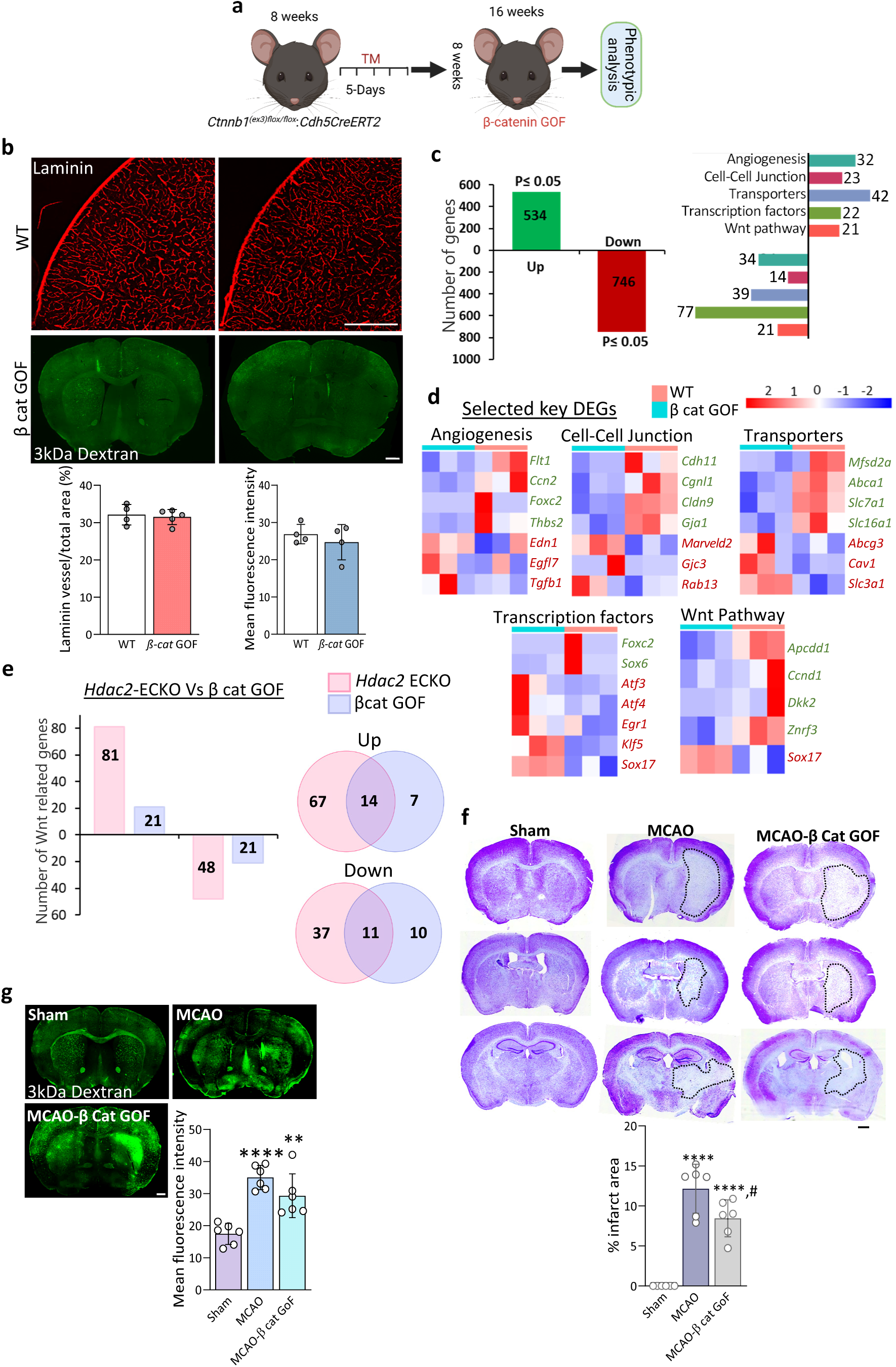
The β-catenin gain-of-function in CNS ECs does not promote angiogenesis or provide neuroprotection against stroke in adult mice. **(a)** Diagrammatic representation of experimental approach to generate EC-specific β-catenin gain of function (β-cat GOF) mice. TM was injected into 8-week-old adult *Ctnnb1^(ex3)flox/flox^*:*Cdh5CreERT2* mice for 5 days, and the brain was collected after 8 weeks for analysis. **(b)** Cortical vessels stained with laminin reveal comparable vascular density in β-catenin GOF and WT mice. BBB permeability assay with a 3 kDa FITC-dextran tracer shows no leakage into the brain parenchyma, similar to WT mice. Images were acquired at 10× magnification and merged via tile scanning. Scale bar: 500 µm. (N =4/group and matching sections were compared between 4 independent mice brains from each group) **(c)** Graph depicting DEGs (p ≤ 0.05 N=3/group) in cortical ECs of β-catenin GOF mice versus WT mice, with upregulated and downregulated genes classified across five key EC functions (angiogenesis, cell-cell junctions, transporters, transcription factors, Wnt pathway). **(d)** Heatmap of selected differentially expressed genes across five key EC functions in β-cat GOF mice versus WT mice. **(e)** Graph showing the number of Wnt-related genes in *Hdac2* ECKO and β-cat GOF mice, along with the overlap in Wnt-related gene expression between the two groups. **(f)** Representative Nissl-stained brain sections from mice subjected to sham surgery, MCAO, and MCAO with β-cat GOF, with infarct areas outlined by black dotted lines. Quantification reveals significantly larger infarct areas in MCAO and MCAO-β-catenin GOF mice compared to sham (****p ≤ 0.0001 vs. sham; #p ≤ 0.05 vs. MCAO; N=6/group, matching sections were compared between 6 independent mice brains from each group). Images were acquired at 10× magnification and merged via tile scanning. Scale bar 500 µm. **(g)** BBB permeability assay using 3kDa tracer showing leakage into ipsilateral brain parenchyma in MCAO and MCAO with β-catenin GOF mice and the fluorescence intensity in brain parenchyma of WT-MCAO and MCAO with MCAO with β-cat GOF mice didn’t show any significant difference. 10 x images are acquired and merged using tile scanning. Scale bar= 500 µm. (****p ≤ 0.0001 vs sham, **p ≤ 0.01 vs sham. N=6/group and matching sections were compared between 6 independent mice brains from each group). All data are presented as mean ± SD.

To investigate whether Wnt activation via β-cat GOF in adult ECs mitigates stroke injury, we induced transient MCAO in mice and administered TM one-hour post-stroke to stabilize β-catenin, continuing treatment for five days. By day 7, Nissl staining showed no reduction in infarct size, and BBB leakage, assessed using 3 kDa dextran, remained pronounced in β-cat GOF mice, comparable to MCAO controls. Post-stroke behavioral analyses also revealed no significant recovery in β-cat GOF mice (Fig. 5f, 5g; Fig. 5, Supplement 5c), indicating that Wnt activation via β-cat GOF alone is insufficient to reduce stroke-induced brain damage or restore BBB integrity.

### β-catenin GOF accelerates angiogenesis in the *Hdac2* ECKO brain

To determine whether Wnt pathway reactivation mediates *Hdac2* ECKO-induced angiogenesis or if further Wnt activation enhances this response, we generated double mutant mice combining *Hdac2* ECKO with β-cat GOF. Eight-week-old mice received five doses of TM, and brains were collected four weeks later (Fig. 6a). IB4 staining revealed significantly increased vessel density in the double mutant compared to both WT and *Hdac2* ECKO mice (Fig. 6b), indicating that β-catenin stabilization augments *Hdac2* deletion-induced angiogenesis. Further, BBB permeability assays showed no significant differences in barrier integrity among the groups (Fig. 6b).

**Figure 6:**
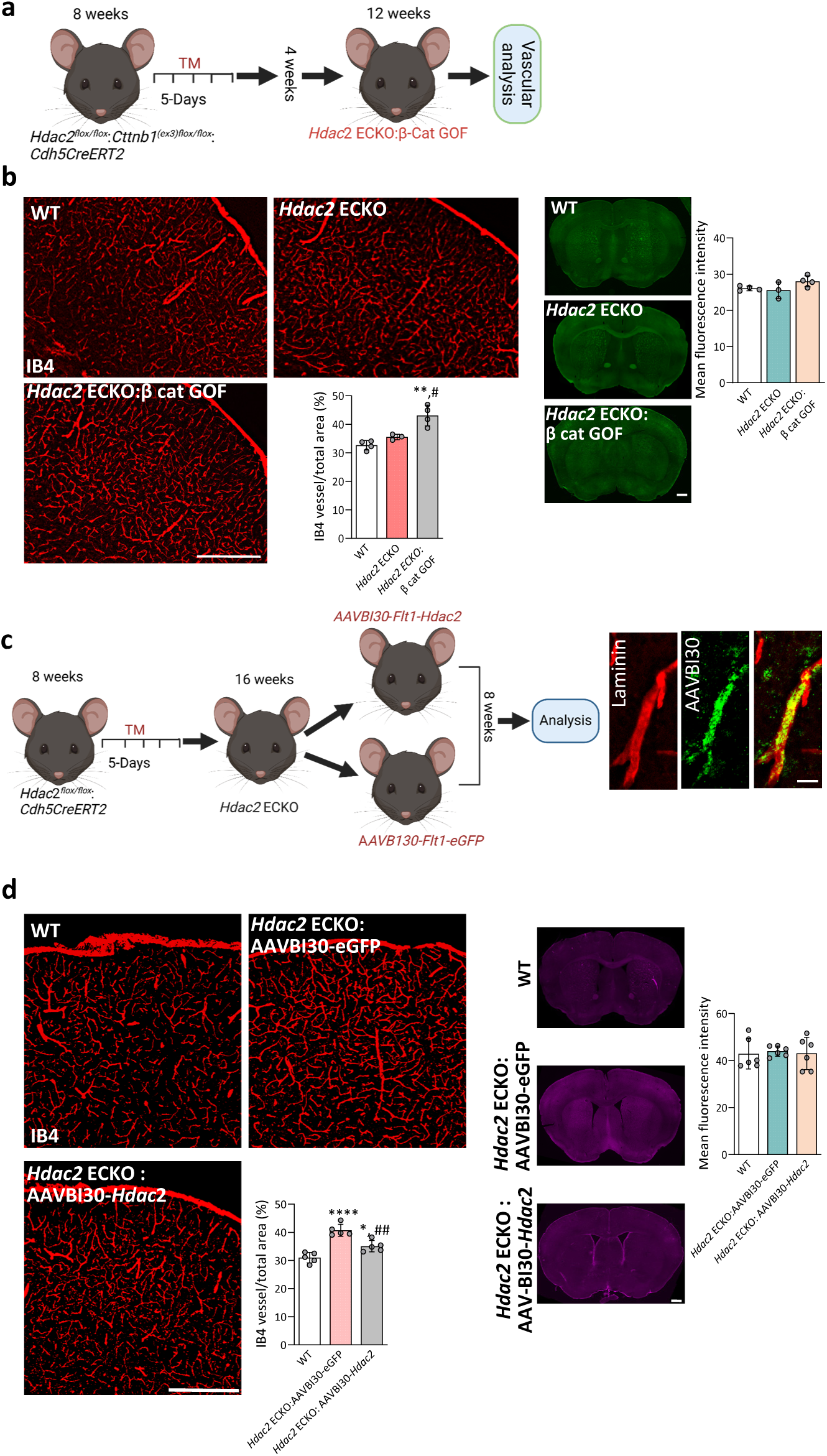
β-catenin GOF accelerates angiogenesis in the *Hdac2* ECKO brain while reintroducing *Hdac2* stabilizes the new vessels. **(a)** Illustration of study methodology. *Hdac2* ECKO;β-catenin GOF double-mutant mice were generated by administering tamoxifen (75 mg/kg body weight) to 8-week-old *Hdac2^flox/flox^*;*Ctnnb1^(ex3)flox/flox^*;*Cdh5CreERT2* mice daily for 5 days. Brains were harvested 4 weeks later for analysis. **(b)** isolectin B4 (IB4) staining and quantification demonstrating increased vascular density in *Hdac2* ECKO:β-catenin GOF double mutant mice after one month compared to WT mice and one month *Hdac2* ECKO mice. Representative images of BBB permeability assay using a 3 kDa-FITC tracer showed no significant BBB leakage in all groups. (**p ≤ 0.01 vs. WT #p ≤ 0.05 vs. *Hdac2* ECKO. Matching sections were analyzed from independent mice from each group. WT N=4/group, *Hdac2* ECKO N=3/group, and double mutant N=4/group). 10 x images are acquired and merged using tile scanning. Scale bar 500 µm. **(c)** Schematic of the experimental framework. Tamoxifen was administered to 8-week-old *Hdac2^flox/flox^*;*Cdh5CreERT*2 mice to generate *Hdac2* ECKO mice. At 8 weeks post-TM, mice were split into two groups: one injected with AAVBI30*-Flt1-eGFP*, the other with AAVBI30*-Flt1-*m*-Hdac2-eGFP*. Mice were sacrificed 8 weeks later, and samples were collected for analysis. Images show brain vessels stained with laminin (vessel marker) and GFP (viral expression), with merged images confirming viral vector expression primarily in brain vessels. Scale bar: 1 µm. **(d)** Representative images of IB4 staining and BBB permeability assay using 3 kDa-Cy3 tracer. Quantification of cortical brain vessels revealed significantly increased vessel density in *Hdac2* ECKO mice with control vector and *Hdac2* ECKO mice with *Hdac2* re-expression vector compared to WT (*p ≤ 0.05, ****p ≤ 0.0001), though the *Hdac2* re-expression group showed significantly reduced density versus the control vector group (##p ≤ 0.01). BBB leakage quantification showed no significant differences between groups. (N= 5/group for IB4; N=6/group for BBB assay. Matching sections were analyszed from independent mice from each group). Images were acquired at 10× magnification and merged via tile scanning. Scale bar: 500 µm. All data are presented as mean ± SD.

### Reintroducing *Hdac2* provides a stable vessel in the *Hdac2* ECKO brain

Since permanent Hdac2 deletion is not physiologically relevant, we tested whether re-expressing Hdac2 in CNS vessels stabilizes newly formed vasculature. We used AAVBI30, which selectively infects the brain microvasculature^17^, to deliver *Hdac2* via an EC-specific AAVBI30-*Flt1*-m-*Hdac2*-eGFP construct; AAVBI30-*Flt1*-m-eGFP served as a control. Two months after TM-induced *Hdac2* deletion, *Hdac2* ECKO mice received intravenous AAVs, and brains were analyzed two months later (Fig. 6c). IB4 staining showed increased vessel density in both AAV-treated groups compared to WT. However, *Hdac2* re-expression significantly reduced vessel density compared to the control virus group (Fig. 6d), indicating suppression of further angiogenesis. BBB integrity, assessed by 3 kDa Cy3-dextran, was preserved in both groups with no tracer leakage (Fig. 6d). These results support the use of transient *Hdac2* suppression to induce angiogenesis.

### *Hdac2* ECKO promotes revascularization and supports structural integrity in the stroke brain

Neuronal loss in the ischemic core is largely complete by day 7 post-stroke, followed by glial scar formation and tissue degeneration^18–20^. Although angiogenesis begins by day 3 and becomes significant by one week, many newly formed vessels undergo apoptosis by day 14^21^, with prominent tissue degradation by one-month post-MCAO. To evaluate whether *Hdac2* ECKO promotes angiogenesis and revascularizes infarcted tissue, we administered TM on day 7 post-stroke, when inflammation subsides, angiogenic signaling remains active, and neuronal death is complete, enabling the study of potential regeneration.

*Hdac2^flox/flox^*:*Cdh5CreERT2* and *Hdac2^flox/flox^*mice underwent MCAO, followed by TM administration on day 7. Brains were collected 8 weeks later; a time point at which *Hdac2* ECKO mice show elevated vascular density. β-cat GOF mice were included for comparison (Fig. 7a). MCAO controls exhibited significant tissue loss in the infarcted hemisphere (Fig. 7b), whereas *Hdac2* ECKO mice showed a markedly improved ipsilateral-to-contralateral tissue ratio, indicating better tissue preservation. This effect was not observed in β-cat GOF mice, confirming the specificity of *Hdac2* ECKO. NeuN staining indicated the presence of neurons in the infarcted hemisphere of *Hdac2* ECKO mice, with a higher ipsilateral-to-contralateral neuron ratio compared to both MCAO and β-cat GOF controls (Fig. 7b). Given the variability in infarct size of MCAO mice, T2-weighted MRI was performed in both MCAO and *Hdac2* ECKO mice at baseline, 7 days post-stroke, and 8 weeks post-treatment. Infarct volume was quantified initially, and ipsilateral cortical volume was measured at 7 days and 8 weeks to assess stroke-induced tissue loss. *Hdac2* ECKO mice showed significantly greater cortical volume preservation compared to MCAO controls, confirming the Nissl and Neun staining results (Fig. 7c; Fig. 7 – figure supplement 7a).

**Figure 7:**
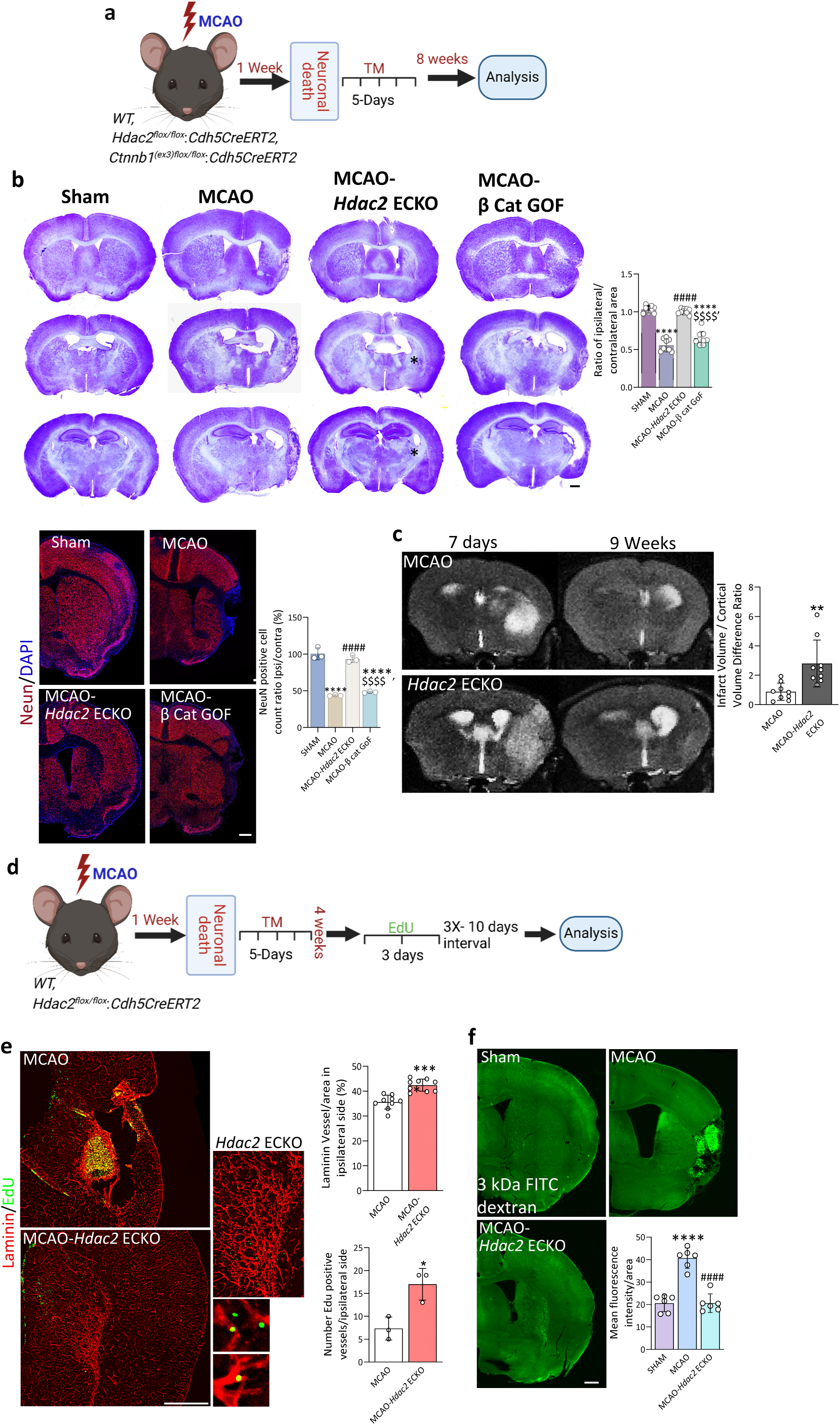
In a 7-day-old stroke-injured brain, *Hdac2* deletion enhances vascularization, prevents tissue degeneration, and promotes neuroregeneration. **(a)** Illustration of the research approach. 8-week-old WT, *Hdac2^flox/flox^*;*Cdh5CreERT2*, and *Ctnnb1^(ex3)flox/flox^*;*Cdh5CreERT2* mice underwent MCAO. One week post-MCAO, TM was administered daily for 5 days. 1 month after MCAO animals received 300 µg of EdU per injection over three consecutive days, with each series separated by a 10-day interval. This treatment regimen was repeated three times, and the mice were sacrificed the day after the final injection for analysis. **(b)** Nissl and NeuN-stained brain sections from sham, MCAO, MCAO-*Hdac2* ECKO, and MCAO-β-cat GOF mice, with infarct areas marked with asterisk in Nissl-stained images. Nine weeks post-MCAO, Nissl staining revealed significant ipsilateral hemisphere loss in MCAO and MCAO-β-cat GOF mice compared to sham, as quantified by the ipsilateral/contralateral area ratio. NeuN immunostaining corroborated this loss, showing a reduced ratio of NeuN^+^ cells in the ipsilateral/contralateral hemispheres. In contrast, MCAO-*Hdac2* ECKO mice exhibited a significant increase in both the Nissl-stained ipsilateral/contralateral area ratio and the NeuN^+^ cell ratio.(****p ≤ 0.0001 vs sham, ####p ≤ 0.0001 vs MCAO, $$$$p ≤ 0.0001 vs MCAO-*Hdac2*-ECKO. N=3/group, matching sections form 3 independent brain) 10 x images are acquired and merged using tile scanning. Scale bar 500 µm. **(c)** Representative immunofluorescence images of laminin and Edu (green fluorescent dots) stained brain sections from MCAO and MCAO-*Hdac2* ECKO mice. A digitally magnified view highlights Edu positive proliferating cells inside vessels within the core infarct region of MCAO-*Hdac2* ECKO mice. Quantification of vascular density and Edu positive vessels in the ipsilateral hemisphere revealed significantly increased vessel density and proliferating vessels (*p ≤ 0.05, ****p ≤ 0.0001 vs MCAO; MCAO, N=3/group, and images were quantified from similar areas of matching sections from three independent brains) in MCAO-*Hdac2* ECKO mice compared to MCAO controls. **(d)** BBB permeability assay using a 3 kDa FITC-conjugated tracer in Sham, MCAO, and MCAO-*Hdac2* ECKO mice. Sham mice exhibited intact vessels with no detectable leakage, whereas MCAO brains showed significant tracer leakage in the infarct core. In contrast, MCAO-*Hdac2* ECKO mice demonstrated no significant leakage. (****p ≤ 0.0001 vs. Sham; ####p ≤ 0.0001 vs. MCAO-*Hdac2* ECKO; N=3/group, two matching sections were taken for analysis from 3 independent brains). Images were acquired at 10× magnification and merged using tile scanning. Scale bar = 500 µm. **(e)** Representative T2-weighted magnetic resonance imaging (MRI) of MCAO and MCAO-*Hdac2* mice. Images depict MCAO and MCAO-*Hdac2* mice at 7 days and 9 weeks post-surgery. The ratio of infarct volume at 7 days to the change in ipsilateral cortical volume between 7 days and 9 weeks was quantified in MCAO and MCAO-*Hdac2* ECKO mice. Compared to MCAO, MCAO-*Hdac2* ECKO brains exhibited a significantly higher ratio (*p ≤ 0.01 vs. MCAO; N = 3 per group; three matching sections per mouse were analyzed). All data are presented as mean ± SD.

To assess *Hdac2* ECKO-induced angiogenesis in the stroke brain, we administered EdU for three consecutive days, followed by a 10-day interval, repeating this cycle three times, starting four weeks after TM treatment. Laminin staining revealed vessel remodeling in the infarct core of *Hdac2* ECKO mice, while MCAO brains showed remodeling limited to the peri-infarct region, with infarct tissue showing no signs of vessels. EdU-laminin co-staining was significantly higher in *Hdac2* ECKO mice, indicating enhanced angiogenesis. Although MCAO brains also showed ongoing angiogenesis, it was less robust. To evaluate BBB integrity, we performed 3 kDa dextran leakage assays. MCAO mice displayed a leaky BBB in the infarct core, whereas *Hdac2* ECKO mice maintained an intact and functional BBB, suggesting restored perfusion in the affected area (Fig. 7d).

### *Hdac2* ECKO promotes post-stroke neurogenesis and neuronal differentiation, facilitating regeneration of infarcted brain tissue

To investigate the mechanisms underlying *Hdac2* ECKO-mediated neuronal regeneration in stroke-damaged brains, we subjected mice to MCAO and induced *Hdac2* deletion as described in Figure 7a. Beginning four weeks after TM injection, proliferating cells were labeled by administering EdU for three consecutive days, followed by a 10-day interval; this cycle was repeated three times (Fig. 8a).

**Figure 8:**
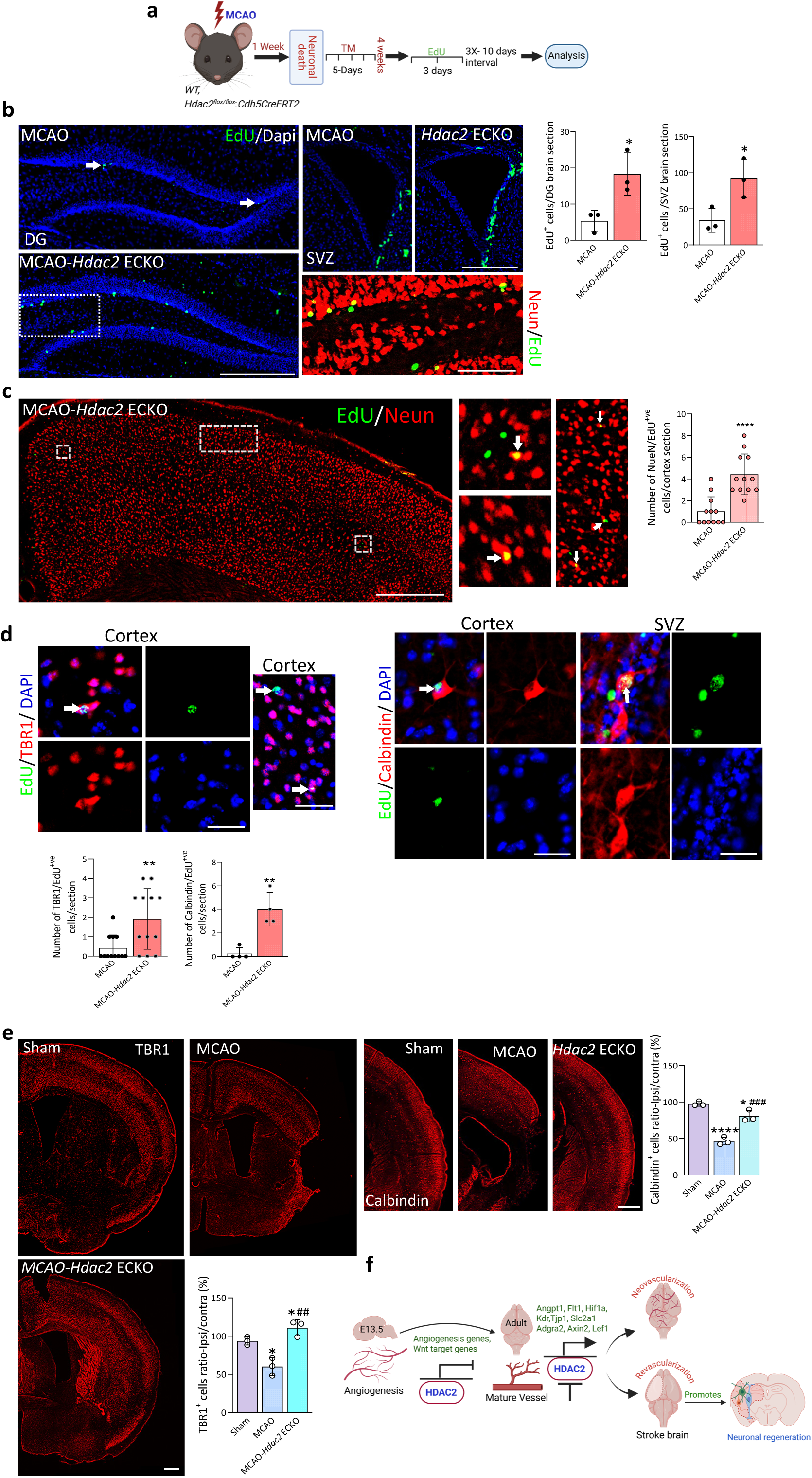
Deletion of CNS EC *Hdac2* induces neurogenesis in the stroke brain. **(a)** Schematic representation of the experimental design. MCAO was induced in 8-week-old *Hdac2^flox/flox^*: Cdh5CreERT2 mice, followed by TM injection for five days starting one week post-MCAO to delete *Hdac2*. The experimental animals received 300 µg of EdU per injection over three consecutive days, with each series separated by a 10-day interval. This treatment regimen was repeated three times, and the mice were sacrificed the day after the final injection. **(b)** Representative images of EdU^+^ cells (green fluorescent dots) in the dentate gyrus and subventricular zone, with quantification showing a significant increase in EdU^+^ cells in MCAO-*Hdac2* ECKO mice compared to MCAO (*p ≤ 0.05 vs. MCAO; MCAO N=3, MCAO-*Hdac2* ECKO N=3). Scale bar: 500 µm. **(c)** Deletion of CNS EC *Hdac2* after MCAO increased the number of proliferating neurons in the cortex. Immunofluorescent image of MCAO-*Hdac2* ECKO cortex stained with NeuN and EdU. Areas within white dotted lines are digitally magnified to show the colocalization of EdU and NeuN. The number of EdU^+^/NeuN^+^ cells in the cortical sections was quantified, showing a significant increase in MCAO-*Hdac2* ECKO compared to MCAO (***p ≤ 0.001 vs. MCAO; N =3 /group; four matching sections per mouse were analyzed). 10x images were acquired and merged using tile scanning. Scale bar: 500 µm. **(d)** 40x images showing EdU^+^ excitatory neurons (TBR1) in the cortex and SVZ (**p ≤ 0.01 vs. MCAO; N = 3/group; four matching sections per mouse were analyzed) and inhibitory neurons (Calbindin) in the cortex and SVZ (**p ≤ 0.01 vs. MCAO; N = 3/group; four matching sections were analyzed) of MCAO-*Hdac2* ECKO. EdU^+^ (green), TBR1^+^ (red), Calbindin^+^ (red), and DAPI^+^ (blue). Scale bar: 5 µm. **(e)** Representative images of TBR1 and Calbindin stained sections in sham, MCAO, and MCAO-*Hdac2* ECKO. The ratio of TBR1^+^ and Calbindin^+^ cells in the ipsilateral versus contralateral cortex was calculated, revealing a significant decrease in MCAO brains compared to sham, whereas MCAO-*Hdac*2 ECKO brains exhibited a significant increase compared to MCAO. Additionally, TBR1^+^ cells in MCAO-*Hdac2* ECKO brains showed a significant increase relative to sham. (*p ≤ 0.05, ****p ≤ 0.0001 vs. sham; ##p ≤ 0.01, ###p ≤ 0.001 vs. MCAO; N=3/group. Matching sections from each group were taken for analysis). 10x images were acquired and merged using tile scanning. Scale bar: 500 µm. All data are presented as mean ± SD.

EdU staining showed increased proliferation in the subventricular zone (SVZ) and dentate gyrus (DG) of *Hdac2* ECKO mice compared to MCAO controls (Fig. 8b). Co-labeling with NeuN confirmed that a substantial portion of EdU⁺ cells differentiated into mature neurons (Fig. 8b). EdU/NeuN co-labeling also confirmed enhanced neurogenesis in the cortex (Fig. 8c). Further analysis showed significantly increased EdU⁺/TBR1⁺ excitatory neurons in layers 5 and 6 of the cortex, and EdU⁺/Calbindin⁺ inhibitory neurons in both the cortex and SVZ of *Hdac2* ECKO mice (Fig. 8d). Confirming the enhanced neurogenesis, quantification of TBR1⁺ and Calbindin⁺ neurons revealed a significant increase in the ipsilateral hemisphere of MCAO-*Hdac2* ECKO mice compared to MCAO controls. In contrast, MCAO mice showed a significant loss of these neuronal populations compared to sham mice (Fig. 8e).

Finally, we examined expression of doublecortin (DCX), a microtubule-associated protein expressed in migrating neuroblasts and immature neurons, widely used as a marker of adult neurogenesis^22–24^. At two months post-injury, DCX⁺ cells were detected in the SVZ of ipsilateral MCAO brains migrating toward peri-infarct region, with a markedly higher density observed in MCAO-*Hdac2* ECKO mice compared to MCAO controls (Fig. 8 - figure supplement 8a). The majority of DCX⁺ cells exhibited morphologies consistent with migrating neuroblasts. Furthermore, MCAO-*Hdac2* ECKO mice exhibited increased cortical thickness in the ipsilateral hemisphere, as determined by comparison with the contralateral side (Fig. 8 – figure supplement 8b), suggesting an enhanced cortical regeneration.

## Discussion

Our study revealed several significant findings. For the first time, we demonstrate epigenetically induced angiogenesis in the adult brain by reactivating the EC developmental program. Importantly, these newly formed vessels are stable, functional, and maintain BBB integrity. We also reveal the epigenetic silencing of developmental Wnt target genes in adult CNS ECs, and show that Wnt pathway activation can further accelerate *Hdac2* ECKO-induced angiogenesis. Additionally, we show that early deletion of *Hdac2* or *Ezh2* in CNS ECs can mitigate stroke-induced transcriptomic alterations and provide neuroprotection. Most critically, we establish that *Hdac2* deletion in CNS ECs promotes revascularization of the stroke-affected brain and supports robust neuronal regeneration within the infarct, suggesting promising therapeutic opportunities for brain angiogenesis and neuronal regeneration.

Brain angiogenesis, which is active during development, later comes under tight regulatory control. Mechanistically, we discovered that epigenetic regulators HDAC2 and PRC2 suppress angiogenic gene expression to promote vascular maturation. Our strategy was to inhibit this repression to reactivate angiogenic programs. EC-specific deletion of *Hdac2* led to the formation of stable, non-hemorrhagic pial and parenchymal vessels, confirmed by vascular staining and perfusion with fluorescently conjugated 3 kDa dextran. Angiogenesis was further validated using the gold-standard proliferation marker EdU. These new vessels exhibited enhanced tight junction (TJ) protein expression, with elevated ZO-1 and CLDN5 levels in proliferating cells, and normal pericyte coverage. Additionally, increased ALDH, GFAP, and AQP4 staining indicated a corresponding rise in astrocyte density, proportional to the increased vessel density in *Hdac2* ECKO mice compared to wild-type controls. Mechanistically, *Hdac2* ECKO activated key angiogenic regulators (*Angpt1*, *Hif1α*, *Kdr*)^25–28^, canonical Wnt targets (*Axin2*, *Lef1*, *Apcdd1*), and core Wnt pathway components (*Adgra2*, *Fzd4*, *Ctnnb1*), ^29–35^all of which are shown essential for brain angiogenesis. Further activation of Wnt signaling via β-cat GOF enhanced this angiogenic response, supporting a functional role for Wnt signaling downstream of HDAC2. Finally, we demonstrate that temporary inactivation of *Hdac2* is feasible: re-expression of *Hdac2* in CNS vessels did not compromise the stability of newly formed vessels, underscoring the therapeutic potential of targeting *Hdac2* for stable adult brain angiogenesis.

Conversely, loss of Ezh2 was associated with vascular regression and increased BBB permeability. Although Ezh2 deletion upregulated several angiogenic (*Notch4*, *Angpt1*), BBB-related (*Cdh5*, *Tjp1*, *Abca1*), transcription factor (*Zic3*), and Wnt target (*Axin2*) genes, the number of regulated genes was substantially lower compared to *Hdac2* deletion, additionally both deletions yielded largely distinct transcriptomic profiles, suggesting that HDAC2 and EZH2 regulate cerebrovascular maintenance through complementary but non-redundant epigenetic mechanisms.

Ischemic stroke causes widespread cerebrovascular damage and persistent transcriptomic alterations in brain ECs^36,37^. Our study demonstrates that immediate deletion of *Hdac2* or *Ezh2* confers significant neurovascular protection, evidenced by improved behavioral performance, reduced infarct volume, and preserved BBB integrity. Transcriptomic analysis revealed extensive dysregulation of EC gene expression 7 days post-MCAO, with upregulation of inflammatory and apoptotic genes and downregulation of pathways essential for vascular stability, including Wnt signaling. In contrast, deletion of *Hdac2*/*Ezh2* markedly reduced MCAO-induced transcriptomic alteration. Comparison of the contralateral hemisphere of sham and *Hdac2* or *Ezh2* deletion showed significantly upregulated genes encoding transcription factors, angiogenesis, and BBB. This epigenetic reprogramming likely underlies the observed protection against ischemic injury. However, our study does not distinguish direct enzymatic targets of HDAC2/PRC2 from downstream, indirect transcriptional changes.

Although post-stroke neurogenesis has been previously reported^38–40^, most studies indicate that newly generated cortical neurons undergo apoptosis shortly after birth^41,42^. While one study showed that SVZ-derived neurons can survive up to 35 days post-stroke^43^, it is well established, including in our current findings, that the infarct core ultimately degenerates into a cavity devoid of structural integrity. *Hdac2* was deleted seven days post-stroke, a time point when inflammation subsides, angiogenic signaling persists, and neuronal death is largely complete^19,20^. Two months after *Hdac2* deletion, stroke brains exhibited substantial tissue preservation compared to MCAO controls, as confirmed by Nissl staining, NeuN immunolabeling, and T2-weighted MRI. EdU-based proliferation assays revealed robust revascularization of the infarct core, indicated by increased EdU⁺/laminin⁺ vessels and elevated vessel density relative to MCAO. Laminin staining further revealed an active vascular remodeling in the infarct zone of Hdac2 ECKO. BBB permeability assays demonstrated persistent leakage in the infarct core of MCAO brains even at two months post-stroke, which was markedly reduced in *Hdac2* ECKO mice.

At the mechanistic level, *Hdac2* deletion enhanced post-stroke neurogenesis. EdU⁺/NeuN⁺ and DCX⁺ cells indicated sustained proliferation and migration of immature neurons toward the infarct core in the MCAO brain. While previous studies have shown limited survival of these cells^40^, *Hdac2* ECKO mice exhibited a marked increase in immature DCX⁺ neurons within the infarct, and the staining pattern suggests their migration towards the infarct core. Moreover, increased co-labeling of EdU with TBR1 and Calbindin, along with higher total numbers of TBR1⁺ excitatory and Calbindin⁺ inhibitory neurons, points to enhanced differentiation and maturation of newborn neurons in the *Hdac2* ECKO stroke brain. Together, our results suggest that *Hdac2* ECKO has the potential to revascularize the stroke brain and promote neuronal regeneration. A limitation of our study is that, although we observed neuronal regeneration, we did not assess the functional integration or activity of the newly generated neurons.

Our data also provide key insights into Wnt pathway regulation in CNS ECs. While Wnt activation during embryogenesis drives robust angiogenesis^8^, this response is absent in adult CNS ECs. Transcriptomic analysis of β-cat GOF mutants revealed activation of only a limited Wnt target, *Apcdd1*, with canonical targets such as *Axin2* and *Lef1* remaining suppressed. β-catenin GOF alone failed to confer neuroprotection or enhance vascular integrity following MCAO. However, when combined with *Hdac2* deletion, it significantly accelerated angiogenesis, indicating that HDAC2 mediated the silencing of developmental Wnt target genes. These findings suggest that Wnt target gene responsiveness differs between developmental and adult CNS vasculature and point to epigenetic silencing as a mechanism restricting Wnt-driven angiogenesis in the mature brain.

## Data availability

Sequencing data have been deposited in GEO under accession codes GSE301293, GSE301448.

## Acknowledgment

We thank Ralf Adams (Max Planck Institute for Molecular Biomedicine) and Ondine Cleaver (The University of Texas Southwestern) for providing the C*dh5(PAC)^Creert2^* mice. We also thank Maketo Taketo (Kyoto University Hospital) for providing the *Ctnnb1*^lox(ex3)^ mice. We thank Dritan Agalliu (Columbia University, Department of Neurology) for critically reading the manuscript and providing valuable suggestions. This work was also supported by NIH grant R01 (R01NS121339), R21 (R21NS135176), and UTHealth Houston startup funds to PKT, and Brain Aneurysm Foundation grant (18412) and Texas Alzheimer’s Research and Care Consortium grant (19593) to ST. All illustrations in the figure were created in BioRender licensed to UTHealth Houston.

## Author contributions

PKT conceived and designed the project. ST executed the experimental work, while CC provided critical support with animal experiments, behavioral assays, and blind data analysis. LK contributed to EdU-based animal experiments, AH did the bioinformatics analysis, and DWM, NJA, and SB provided valuable conceptual input.

## Materials and Methods

### Experi*mental Animals*

*Hdac2^flox/flox^* (Strain #:022625)*, Ezh ^flox/flox^* (Strain #:022616), mice of both sexes were purchased from Jackson Laboratories. *Ctnnb1*^(ex3)*flox/flox*^ (Maketo Taketo) was transferred from the Vanderbilt school of medicine, Nashville, Tennessee, USA. Tamoxifen-inducible driver *Cdh5CreERT2* (Ralf Adams) was transferred to the ≤I facility from UT Southwestern. All mice were fed with a standard laboratory diet and water, maintained under standard laboratory conditions (temperature: 25 ± 2°C, humidity: 60 ± 5%, 12 h dark/light cycle) with free access to a standard pellet diet and water ad libitum. Animal procedures were carried out under the oversight of the Animal Care and Use Committee of the University of Texas Health Science Center at Houston (protocol # AWC-22-0108) and in strict compliance with National Institutes of Health guidelines.

### Generation of transgenic mice

For generating transgenic mice we crossed *Hdac2^flox/flox^*or *Ezh2 ^flox/flox^or Ctnnb1*^(ex3)*flox/flox*^ mice with *Cdh5CreERT2* mice. To activate the cre-recombinase (gene deletion), we injected five doses of tamoxifen (Sigma T5648) diluted in corn oil (Sigma C8267) at a dose of (75mg/Kg) for five days.

### Materials

Paraformaldehyde (PFA, P6148), glyoxal (128465), xylene for histology (XX0060-4), Percoll (17-0891-01) were purchased from Sigma-Aldrich (MilliporeSigma, Burlington, MA, USA). Phosphate-buffered saline (PBS) (CA008-050) was obtained from Gen DEPOT, Katy, TX, USA. Permount mounting medium (SP15-500), hydrochloric acid (A481212), and sodium hydroxide (S320-500) were purchased from Fisher Scientific, Waltham, MA, USA. Crystal violet solution (PS102) was procured from FD Neuro Techkologies, Columbia, USA.

### Pial vessel imaging and calculation of pial vessel density

Mice were euthanized according to institutional guidelines, using CO_2_ euthanasia, following. American Veterinary Medical Association Guidelines. 70% (vol/vol) ethanol was sprayed onto the head/neck area and remove the brain was isolated by removing the skull carefully to avoid damaging the brain. Brain was transferred in to 35 mm Petri dish filled with cold 1× PBS on ice and imaged using ZEISS SteREO Discovery. V12 (ZEISS Discovery. V12 steREO, ZEISS Microscopy, Dublin, CA, USA) Density of the pial vessel was calculated using Angiotool^44^ as %vessel detected/area of the brain.

### Determination of BBB permeability and quantification of vessel percentage area in wild-type and mutant brain sections

Adult mice were deeply anesthetized and 50 μl of 3 kDa FITC conjugated dextran was injected into the left ventricle with Hamilton syringe. After 5 minutes of circulation, the right atrium was cut open, and 10 ml of cold PBS, followed by 10 ml 4% PFA (Paraformaldehyde), was injected into the left ventricle using a Hamilton syringe at a rate of 5 ml/minute. The brain was dissected and fixed by immersion in 4% PFA at 4°C overnight, followed by storage in PBS with azide. 30 μm vibratome (semiautomatic vibratome, LEICA VT 1000 S, Leica Biosystems, Deer Park, IL, USA) sections were generated, washed in PBS, mounted using a DAPI Fluoromount-G mounting medium, and imaged using a fluorescent microscope (Leica Thunder imager). Matching sections were taken from WT and mutant brains, examined for BBB leakage, and quantified using Image J software. For comparing vessel density between WT and mutant brains, matching brain sections were taken from both the groups, stained with isolectin B4/Laminin to visualize blood vessels, mounted using a DAPI Fluoromount-G mounting medium and imaged using a fluorescent microscope (Leica Thunder imager). The percentage of the area occupied by vessels in the brains was calculated using Angiotool software.

### Isolation of ECs from mouse brain cortex using flow cytometry (FACS)

For isolating ECs from the mouse brain cortex, we followed the protocol outlined in our previous publication, which is based on the method developed by Crouch and Doetsch ^45^. Briefly, mice were humanely euthanized in accordance with institutional guidelines. Skull was cut opened, the brain tissue was collected into cold 1X PBS. The cortex was isolated after peeling off the pial vessel by rolling the brain over the Whatman filter paper. The cortical brain tissue was chopped and placed into a tissue collection solution on ice; homogenization was done using a Dounce homogenizer. The homogenate was centrifuged at 300g for 5 minutes at 4 °C. The pellet was resuspended in collagenase/dispase solution and incubated for 30 minutes at 37 °C with constant rotation in a hybridization oven. The digested tissue was centrifuged at 300g for 5 minutes at 4°C. After confirming the digestion under a microscope, the pellet was suspended in a trituration solution and triturated by pipetting up and down (100 times). Percoll gradient centrifugation was performed using 22% percoll (vol/vol), at a centrifugation speed of 560g for 10 minutes at 4 °C. The obtained pellet was resuspended in HBSS/BSA/glucose buffer and incubated with antibodies for 20 minutes on ice without agitation (CD31-APC: 551262, 1:50), CD41-PE: 558040, 1:200 and CD45-PE:553081,1:200). Excess unbound antibodies were removed by adding extra HBSS/BSA/glucose buffer and centrifuging at 300g at 4 °C for 5 min. The pellet was resuspended in HBSS/BSA/glucose buffer, filtered using a 40-µm cell strainer, and DAPI (1µg/ml) was used to exclude dead cells during FACS sorting. CD45^+v^e and CD41^+ve^ cells were excluded and live CD31^+ve^ endothelial cells (CD31^+ve^ CD41^−ve^CD45^−ve^) were collected.

### Ultra-low mRNA sequencing

The FACS sorted cells were suspended in solution as suggested by SMART-Seq v4 PLUS Kit User Manual (Takara Bio, Japan). The sorted cells were frozen on dry ice as soon as the sorting was done at The University of Texas Health Science Center in Houston (CPRIT RP180734). Libraries were prepared with SMART-Seq V4 PLUS Kit (R400752, Takara Bio, Japan) following the manufacturer’s instructions. Agilent High Sensitive DNA Kit (#5067-4626) by Agilent Bioanalyzer 2100 (Agilent Technologies, Santa Clara, USA) was used to check the quality of the final libraries and the library concentrations were determined by qPCR using Collibri Library Quantification kit (#A38524500, Thermo Fisher Scientific, Texas, USA). The libraries were pooled evenly and went for the paired-end 150-cycle sequencing on an Illumina NovaSeq X plus System (Illumina, Inc., USA).

### Middle Cerebral Artery Occlusion (MCAO) model of ischemic stroke

Mice were prepared one day ahead for the surgery by removing the hair in the thoracic area. On the day of surgery, mice received buprenorphine (0.003mg/mouse) as analgesic and bupivacaine (2.5mg/Kg body weight) as local analgesic along with Isoflurane to induce anesthesia. Briefly, the right common carotid artery and the right external carotid artery was surgically exposed and ligated following a ventral midline neck incision, and ischemia was induced by inserting a silicone-coated nylon monofilament (Doccol Corp, USA) through the common carotid artery into the internal carotid artery. Regional cerebral blood flow (rCBF) was measured throughout the operation by laser-Doppler flowmetry (Perimed AB, Sweden) with a flexible probe fixed on the skull 1 mm posterior and 6 mm lateral to the bregma. Occlusion was made sure by a decrease of regional cerebral blood flow (rCBF) by at least 80% of the start value. During occlusion, rCBF was maintained below 20% of baseline for 60 minutes, after which the occluding filament was withdrawn to allow reperfusion. Reperfusion was confirmed by the rise in the rCBF above 50% of baseline within 10 min after filament removal. In sham animals, a suture was introduced up to the origin of the middle cerebral artery and then removed immediately, and mice were kept under anesthesia for 60 minutes.

### Nissl staining

Brain sections were loaded onto microscopic slides and were dried at 37°C overnight. On the next day, sections were hydrated for 3 minutes in water and covered with FD Cresyl Violet Solution for 5 minutes. Followed by distaining using 0.1% Glacial acetic acid in 100% ethanol, sections were dehydrated using ascending grades of ethanol (70%, 90%, and 100%), cleared in xylene, and mounted using Permount mounting medium. ImageJ software was used for calculating infarct area and results were expressed as % of infarct area.

### Re-expressing *Hdac2* in CNS ECs of *Hdac2* knock out mice

Eight-week-old mice received five doses of TM (75 mg/kg) to delete *Hdac2*, as mentioned earlier. Eight weeks later, the mice received a retro-orbital injection of AAVBI30-*Flt1*-m-*Hdac2*-*eGFP* at a concentration of 2×10^10^ viral particles per mouse to re-express *Hdac2*. Control cohorts received AAVBI30-*Flt1*-m-*eGFP*. The mice were sacrificed eight weeks later to analyze the vascular phenotype.

### Immunohistochemistry

Brain sections were observed under the bright field microscope (ZEISS Discovery. V12 steREO, ZEISS Microscopy, Dublin, CA, USA) to ensure matching sections were selected for staining. Sections were put in PBST (PBS containing 0.02% triton X 100) for 30 minutes and were then incubated in selected primary antibodies in PBST overnight at 4°C with gentle shaking. The following antibodies were used: laminin (1:200, Anti-Laminin _-2 Antibody (4H8-2): sc-59854; Santa Cruz Biotechnology, Santa Cruz, CA, USA), isolectin B4 (1:200, DL-1207 Vector Laboratories, California, USA), claudin 5 (1:200, 4C3C2; Invitrogen, Waltham, MA, USA), Glial fibrillary acidic protein (GFAP) (1:100, sc-33673, Santa Cruz Biotechnology, Santa Cruz, CA, USA), aquaporin-4 (1:100, 20104, BiCell Scientific, MO, USA), Aldehyde dehydrogenase 1 (Aldh1) (1:50, 85828S, Cell Signaling Technologies, Danvers, MA, USA). NeuN (1:1000, ABN78, MilliporeSigma, Burlington, MA, USA), TBR1 (1:400, (1:400, TBR1(D6C6X, 49661, Cell Signaling Technologies, Danvers, MA, USA), Calretinin (1:400, Calretinin (E7R6O) 92635, Cell Signaling Technologies, Danvers, MA, USA), Calbindin (1:200 Calbindin (D1I4Q), 13176, Cell Signaling Technologies, Danvers, MA, USA) zonula occludens-1 (ZO1) (1:250, 00236; BiCell Scientific, MO, USA and platelet-derived growth factor receptor beta (PDGFRB) (1:200, 31695; Cell Signaling Technologies, Danvers, MA, USA). Next day, the brain sections were washed thrice with PBS and stained with conjugated antibodies (laminin and isolectin B4) were mounted on glass slides using DAPI Fluoromount-G (SouthernBiotech Birmingham, AL, USA) fluorescent microscope (Leica Thunder imager) was used for imaging. Non-conjugated antibodies (CLDN5, AQP4, ZO1, ALDH1, GFAP, PDGFRB, NeuN, TBR1, Calbindin, calretinin) were detected by probing with a biotin-conjugated secondary antibodies from Vector Laboratories in PBST (1:200) for 2 hours. The following secondary antibodies were used: Goat Anti-Rabbit IgG Antibody (H + L), Biotinylated (BA-1000-1.5), Goat Anti-Mouse IgG Antibody (H + L) and Biotinylated (BA-9200-1.5). After incubation, sections were washed with PBS and incubated with Streptavidin, DyLight® 488/594 (SA-5488-1/SA-5549-1), washed with PBS, mounted and imaged using a fluorescent microscope (Leica Thunder imager). Quantification of IB4, Laminin, and Aquaporin positive vessels was done using Angiotool as %vessel detected/area of the brain. The density of ALDH1, GFAP, NeuN, and TBR1 calbindin-positive cells was quantified using ImageJ software.

### *In vivo* EdU labeling detection in mice brains

To detect the vascular proliferation EdU was administered into mice by intraperitoneal injection (300 µg/injection) for three days before the sacrifice. To detect the neuronal proliferation, EdU was administered to mice by intraperitoneal injection (300 µg/injection) for three days for three times at a day interval between each set of injections. The population of EdU^+^ cells in the brain section was detected by following the protocol in the Edu detection kit (Click-iT Plus EdU Alexa Fluor 488 imaging kit, Invitrogen, Thermo Fisher Scientific, Texas, USA). Briefly brain sections were washed with 3% bovine serum albumin in PBS for five minutes (two times) and incubated with 0.5% PBST (PBS containing 0.5% Triton X100) for 30 minutes. After washing with 3% bovine serum albumin in PBS for five minutes (two times), sections were incubated with a reaction cocktail for 30 minutes and washed with 3% bovine serum albumin in PBS and mounted onto the slide and examined and counted for EdU-positive cells as green fluorescent spots.

### Magnetic resonance imaging

In vivo, magnetic resonance imaging (MRI) was performed using a 70/30 7 Tesla small animal MRI scanner (Bruker Biospin MRI, Inc., Billerica, MA) equipped with a 35mm ID 1H transmit/receive volume coil. Animals were anesthetized using 2% isoflurane. Diffusion tensor imaging was used to evaluate the changes of infarction in mice allocated to MCAO (N=3) and MCAO-Hdac2 ECKO (N=3) after 7 days and 2 months of surgery. T2 (RARE) 2D images were acquired in the Coronal and Axial planes using the following parameters: T2 RARE Cor, FOV 40×30 mm, Matrix size = 256×192, spatial resolution of 156 uM, TE=57 ms; RT= 3000 ms, slice thickness, 0.75 mm; slice gap, 0.25mm, RARE factor =12, NEX = 4. T2 RARE Ax, FOV 30×22.5 mm, Matrix size = 256×192, spatial resolution of 117uM, TE =57ms, RT= 3000 ms, slice thickness= 0.75 mm; slice gap, 0.25mm, RARE factor = 12, NEX = 4. Infarction volume was analyzed using ITK-SNAP^46^.

### Behavioral tests

All behavioral tests were conducted in assigned behavioral testing rooms during the light phase of the light/dark cycle in a random order, by observers who did not know the group of the mice until the test had been completed. After each test, the equipment was cleaned with 30% ethanol to eliminate olfactory cues. Activity and behavior of mice were observed using an automatic video tracking system for recording and analysis (Ethovision XT) were used. All mice were brought to the behavior room at least 30 minutes before beginning the testing.

### Open field test (OFT)

The anxiety and general ambulatory capacity of animals were assessed by using OFT. The open field apparatus consists of an arena (made of ¼ inch thick, white material) that is 40 cm×40 cm × 30 cm in size. Mice were placed in the center of the square. The behavior of the animals in the open field was recorded for 20 min. Overall activity in the box (measured with videotrack) was measured as well as the amount of time and distance traveled in the center area of the maze. This paradigm is based on the idea that mice will naturally prefer to be near a protective wall rather than exposed to danger out in the open. Data analysis was performed using the Ethovision XT software.

### Y-maze test

Spatial working memory was assessed using a Y-maze. Mice were placed at the center of the Y-maze. Mice were tested with no previous exposure or habituation to the maze. A spontaneous alternation was defined as an entry into three different arms on consecutive choices. The total distance traveled, a number of entries, and a number of alternations were recorded and analyzed using Ethovision XT software.

### Barnes maze test

Barnes maze is 91.44 cm, having sixteen 1.9 cm holes, which are 1 cm from the edge. One hole (hole 16) has an escape box (7.5 cm × 15 cm × 12.7 cm in height with a metal grid of 1.4 cm spacing) and escape box is in the wooden holder. Visual cues were placed on the wall about the table level. Training is consisted of 4 trials daily for 4 days. Each trial was terminated when the mouse completely escaped through the target hole or after 5 minutes had elapsed. After each trial, the mouse was allowed to remain in the escape chamber for 30 s. A probe trial was tested at 24 h after training. Recall was assessed by comparing the number of entries into each hole around the maze. Acquired data was analyzed using Ethovision XT software.

## STATISTICAL ANALYSIS

Statistical analysis was performed using the GraphPad Prism 6 software. Replicated biological samples were used for statistical analysis as indicated by sample size (n) in figure legends. Results were expressed as mean ± SD (standard deviation). Significance between two groups was calculated using an unpaired Student’s t-test (two-tailed) or paired Student’s t-test for two groups of values representing paired observations. One-way ANOVA was used to assess the statistical significance between multiple-group comparisons, along with appropriate *post hoc* tests (Bonferroni’s or Tukey’s multiple comparison tests). p value ≤ 0.05 was considered as statistically significant.

---

**Figure 1 supplement-1:**
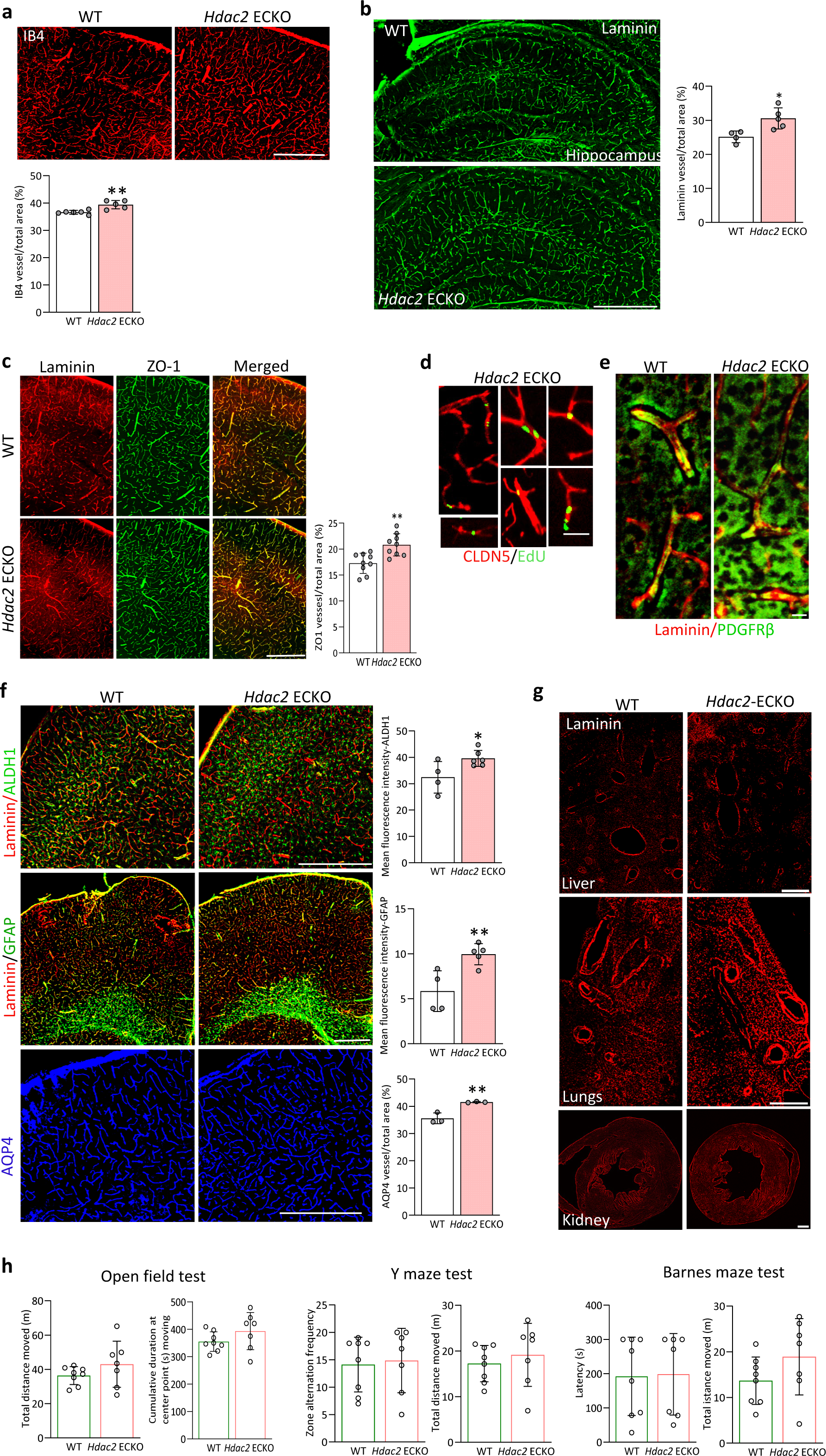
Characterization of CNS vessels in Hdac2 ECKO and behavior analysis of Hdac2 ECKO and Ezh2 ECKO. **(a)** IB4 staining displayed significantly increased vascular density in *Hdac2* ECKO mice cortex compared to WT (**p ≤ 0.01vs WT. WT N=6 and *Hdac2* ECKO N=5 quantifications were performed on matching sections from independent brains). Scale bar: 500µm. **(b)** Laminin staining displayed a significant increase in hippocampal vessel density in *Hdac2* ECKO mice compared to WT (*p ≤ 0.05 vs WT. WT N=4 and *Hdac2* ECKO N=5 quantifications were performed on matching sections from independent brains). 10 x images are acquired and merged using tile scanning. Scale bar 500 µm. **(c)** Co-staining of Laminin and ZO1 displaying significantly increased ZO1 stained vessels in *Hdac2* ECKO mice (**p ≤ 0.01vs WT N=3/group, three similar regions of interest in cortex from matching sections were selected from each mice brains). Scalebar: 500 µm. **(d)** Tight junction protein CLDN5 and EdU co-staining in *Hdac2* ECKO mice vessels indicate newly formed vessels express CLDN5. Scale bar: 50µm. **(e)** Co-staining of laminin for vessel and PDGFRβ for pericytes displaying pericyte covering on vessels of both WT and *Hdac2* ECKO mice. Scale bar: 50µm. **(f)** Representative images of brain sections from WT and *Hdac2* ECKO mice stained for the astrocyte markers ALDH1 and GFAP, co-stained with laminin, and the astrocyte endfeet marker AQP4. Quantification revealed a significant increase in the density of ALDH1 (*p ≤ 0.05 vs. WT; WT: N=4, *Hdac2* ECKO: N=6), GFAP (**p ≤ 0.01 vs. WT; WT: N=4, *Hdac2* ECKO: N=5), and AQP4 (**p ≤ 0.01 vs. WT; WT: N=3, *Hdac2* ECKO: N=3) in *Hdac2* ECKO mice. Images were acquired at 10× magnification and merged using tile scanning. Scale bar: 500 µm. Quantifications were performed on matching sections from independent brains. **(g)** Representative images of laminin staining in liver, lung, and kidney tissues from WT and *Hdac2* ECKO mice. Scale bar: 500 µm. **(h)** Analysis of mice behavior using open field test, Barnes maze test, and Y maze test showed no significant difference in *Hdac2* ECKO mice behavior compared to WT (WT N=8 and *Hdac2* ECKO N=7). All data are presented as mean ± SD.

**Figure 1 supplement-2:**
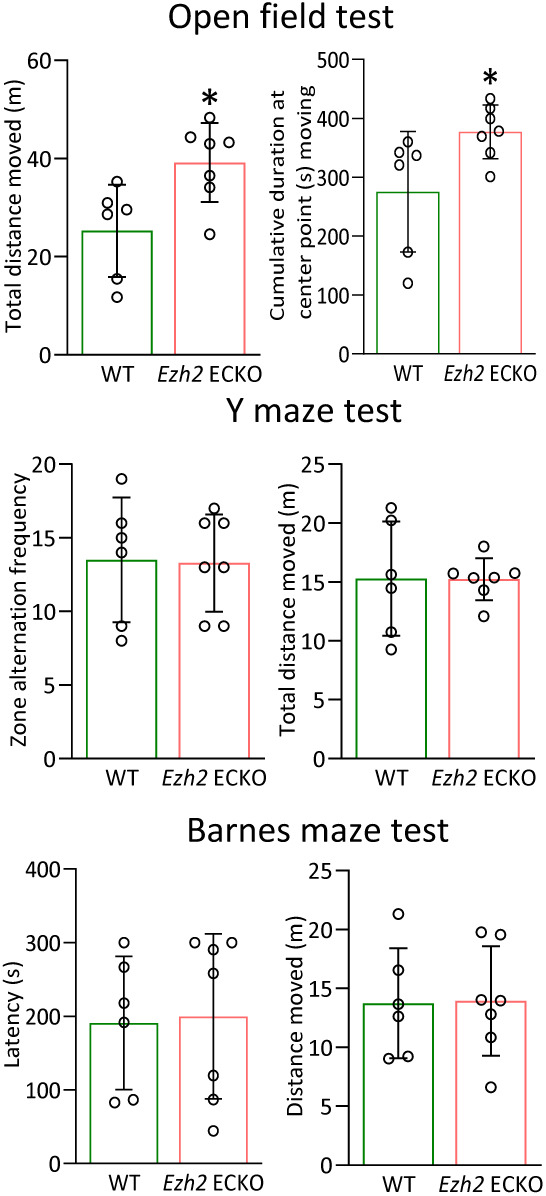
Behavioral abnormalities in *Ezh2* ECKO mice. Behavioral analysis of WT and *Ezh2* ECKO mice using the open field test revealed a significant increase in total distance moved and cumulative duration spent at the center in *Ezh2* ECKO mice compared to WT. Additionally, the Barnes maze and Y-maze tests showed no significant behavioral differences in *Ezh2* ECKO mice. (*p ≤ 0.05 vs. WT; N=6 and *Ezh2* ECKO N=7). All data are presented as mean ± SD.

**Figure 2 Supplement 1:**
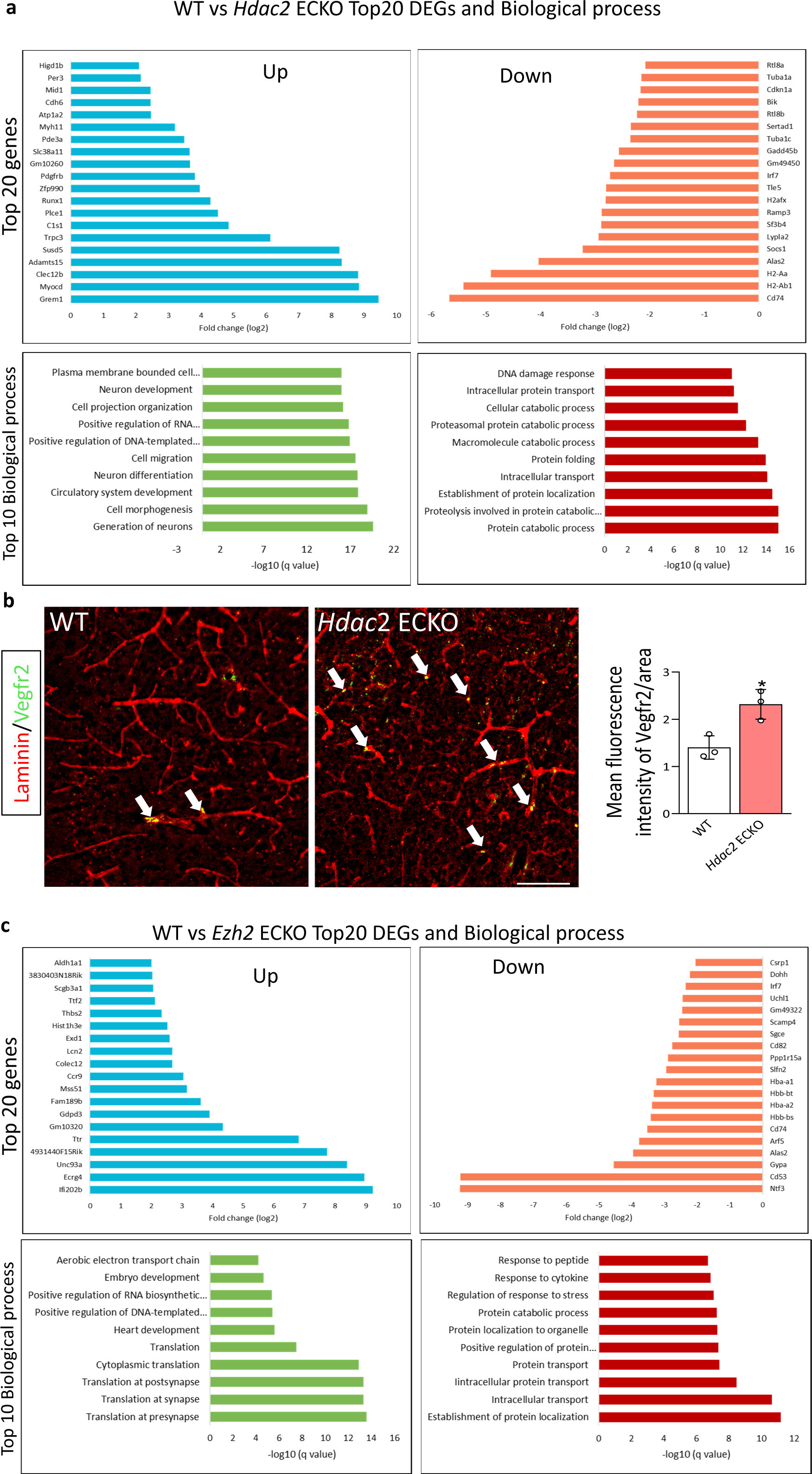
Top 20 upregulated and downregulated DEGs, along with the biological processes associated with DEGs in *Hdac2* ECKO and *Ezh2* ECKO compared to control. **(a)** Diagram depicting Top 20 DEGs (upregulated and downregulated) in *Hdac2* ECKO ECs based on fold change (padj ≤ 0.05) and associated top 10 biological processes (upregulated and downregulated). **(b)** Representative images of brain sections from WT and *Hdac2* ECKO mice co-stained for Vegfr2, an angiogenic marker and laminin. Quantification revealed a significant increase in the density of vegfr2 (*p ≤ 0.05 vs. WT; WT: N=3, *Hdac2* ECKO: N=3) in *Hdac2* ECKO mice. Images were acquired at 10× magnification and merged using tile scanning. Scale bar: 100 µm. Quantifications were performed on matching sections from independent brains. Data is presented as mean ± SD. **(c)** Image showing Top 20 DEGs (upregulated and downregulated) in *Ezh2* ECKO ECs based on fold change (padj ≤ 0.05) and associated top 10 biological processes (upregulated and downregulated).

**Figure 2 supplement 2:**
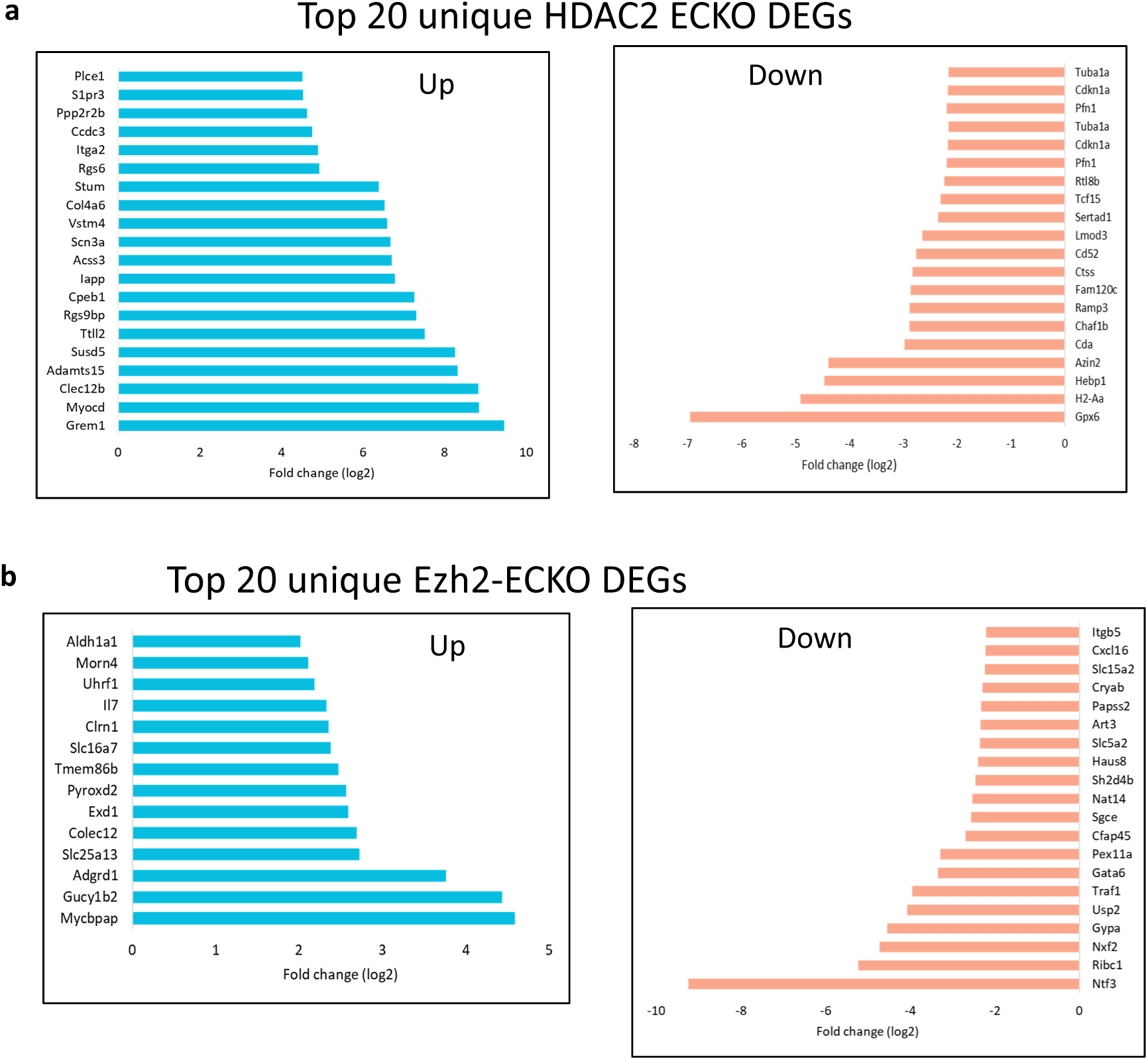
Gene Ontology (GO) enrichment analysis of unique DEGs in *Hdac2* ECKO and *Ezh2* ECKO ECs using ToppGene, highlighting the top five biological processes and disease associations. (**a**) Schematic representation of the top 20 unique genes and the top 10 biological terms associated with upregulated and downregulated genes in *Hdac2* ECKO ECs, based on fold change and a significance threshold of padj ≤ 0.05. (**b**) Schematic representation of the top 20 unique genes and the top 10 biological terms associated with upregulated and downregulated genes in *Ezh2* ECKO ECs, based on fold change and a significance threshold of padj ≤ 0.05.

**Figure 4 supplement:**
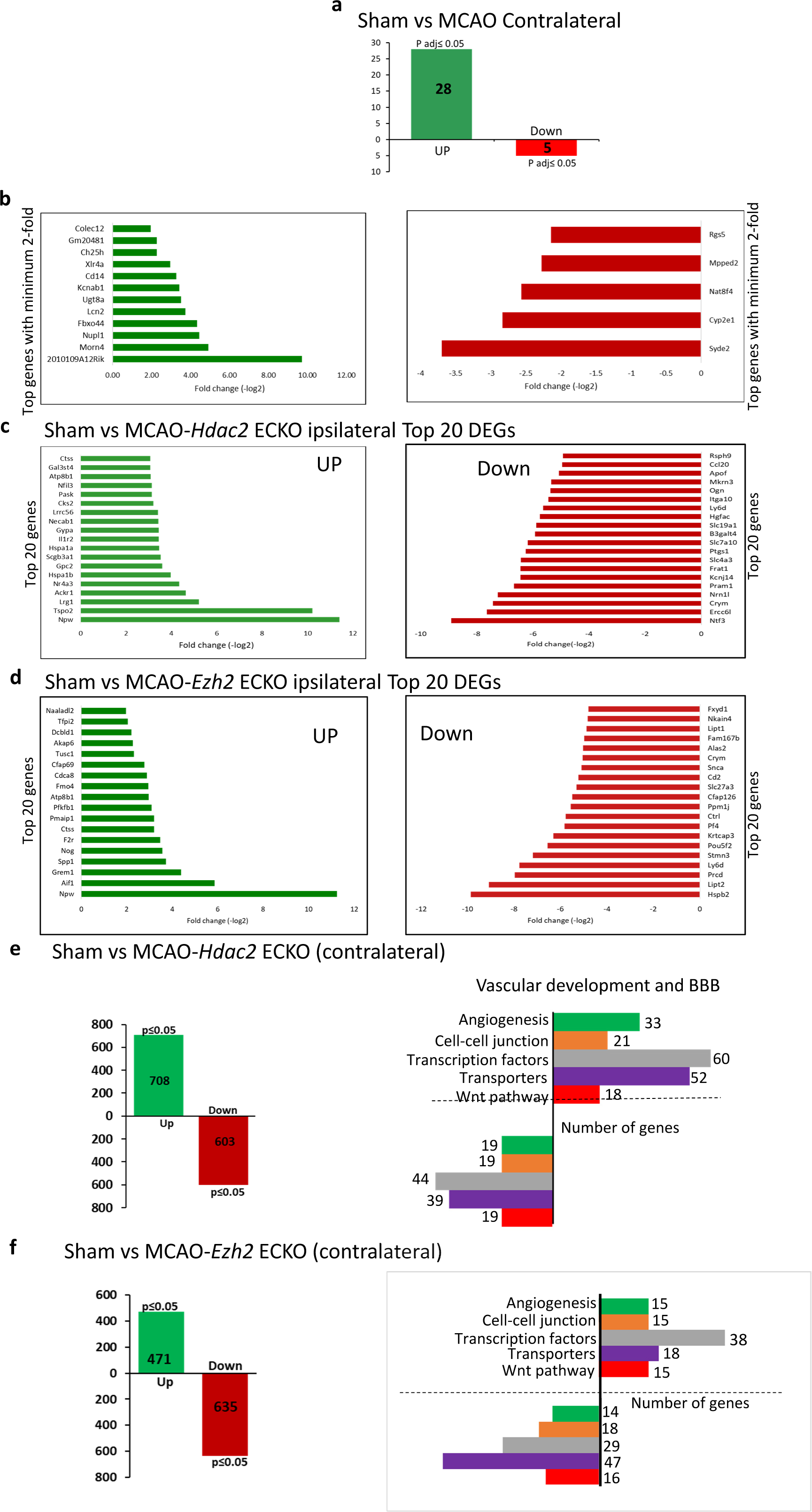
Sham vs. MCAO Contralateral DEGs and Top 20 DEGs, Biological Processes of DEGs in Sham vs. MCAO Ipsilateral, Sham contralateral vs. MCAO-*Hdac2/Ezh2* ECKO contralateral. **(a)** Graph depicting significantly upregulated and downregulated genes (padj ≤ 0.05) in contralateral cortical ECs from sham and WT MCAO mice. **(b)** Image highlighting top upregulated and downregulated DEGs (minimum 2-fold change, padj ≤ 0.05) in contralateral cortical ECs of MCAO mice. **(c)** List of the top 20 DEGs (upregulated and downregulated) in ipsilateral cortical ECs of MCAO-*Hdac2* ECKO mice versus sham, ranked by fold change (pad ≤ 0.05). **(d)** List of the top 20 DEGs (upregulated and downregulated) in ipsilateral cortical ECs of MCAO-*Ezh2* ECKO mice versus sham, ranked by fold change (padj ≤ 0.05). **(e)** Graph depicting significantly upregulated and downregulated genes (p≤ 0.05) in contralateral cortical ECs from sham and MCAO-*Hdac2* ECKO mice and classification of differentially expressed genes (DEGs) into vascular development-related and BBB gene ontology (GO) terms. **(f)** Graph showing the number of significantly upregulated and downregulated genes (p ≤ 0.05) in cortical ECs of in contralateral cortical ECs from sham and MCAO-*Ezh2* ECKO mice mice and classification of differentially expressed genes (DEGs) into vascular development-related and BBB gene ontology (GO) terms.

**Figure 5 Supplementary:**
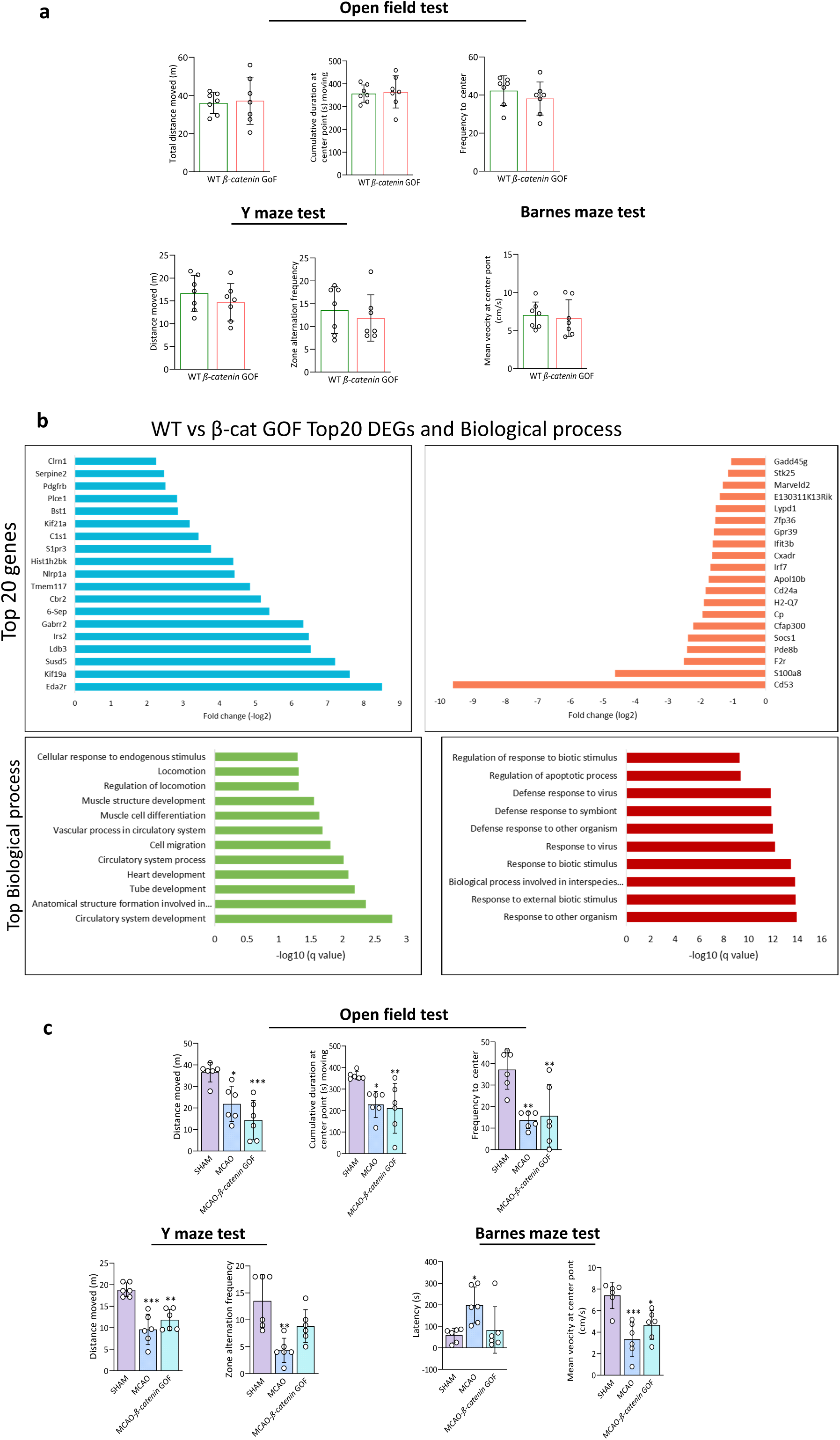
Behavioral Effects, Top 20 Differentially Expressed Genes (DEGs), and Biological Processes in β-Catenin Gain-of-Function. **(a)** β-cat GOF mice behaves similar to that of WT mice. The behavior tests open field test (OFT), Barnes maze test (BMT), and Y maze test (YMT) were conducted in β-cat GOF, and obtained data shows regaining β-cat didn’t change mice behavior significantly compared to WT mice (N=7/group). **(b)** Image displaying top 20 genes that are differentially expressed (upregulated and downregulated) in β-cat GOF mice brain ECs compared to WT mice, based on fold change (p ≤ 0.05) and top 10 biological process associated with upregulated and downregulated genes. **(c)** In the Open Field Test, β-cat GOF did not mitigate deficits in MCAO mice across three parameters: total distance moved, cumulative duration at the center, and frequency of center entries, all significantly reduced versus sham. In the Y Maze Test, MCAO and MCAO-β-cat GOF mice exhibited a significant decrease in total distance moved; however, MCAO-β-cat GOF showed no significant difference in zone alternation compared to sham or MCAO. In the Barnes Maze Test, MCAO mice displayed significantly increased latency to reach the target hole, while MCAO-β-cat GOF showed no significant difference from sham or MCAO. Mean velocity at the center point was significantly reduced in both MCAO and MCAO-β-cat GOF groups compared to sham. (*p ≤ 0.05, **p ≤ 0.01, ***p ≤ 0.001 vs sham N=6/group). All data are presented as mean ± SD.

**Figure-7 Supplement:**
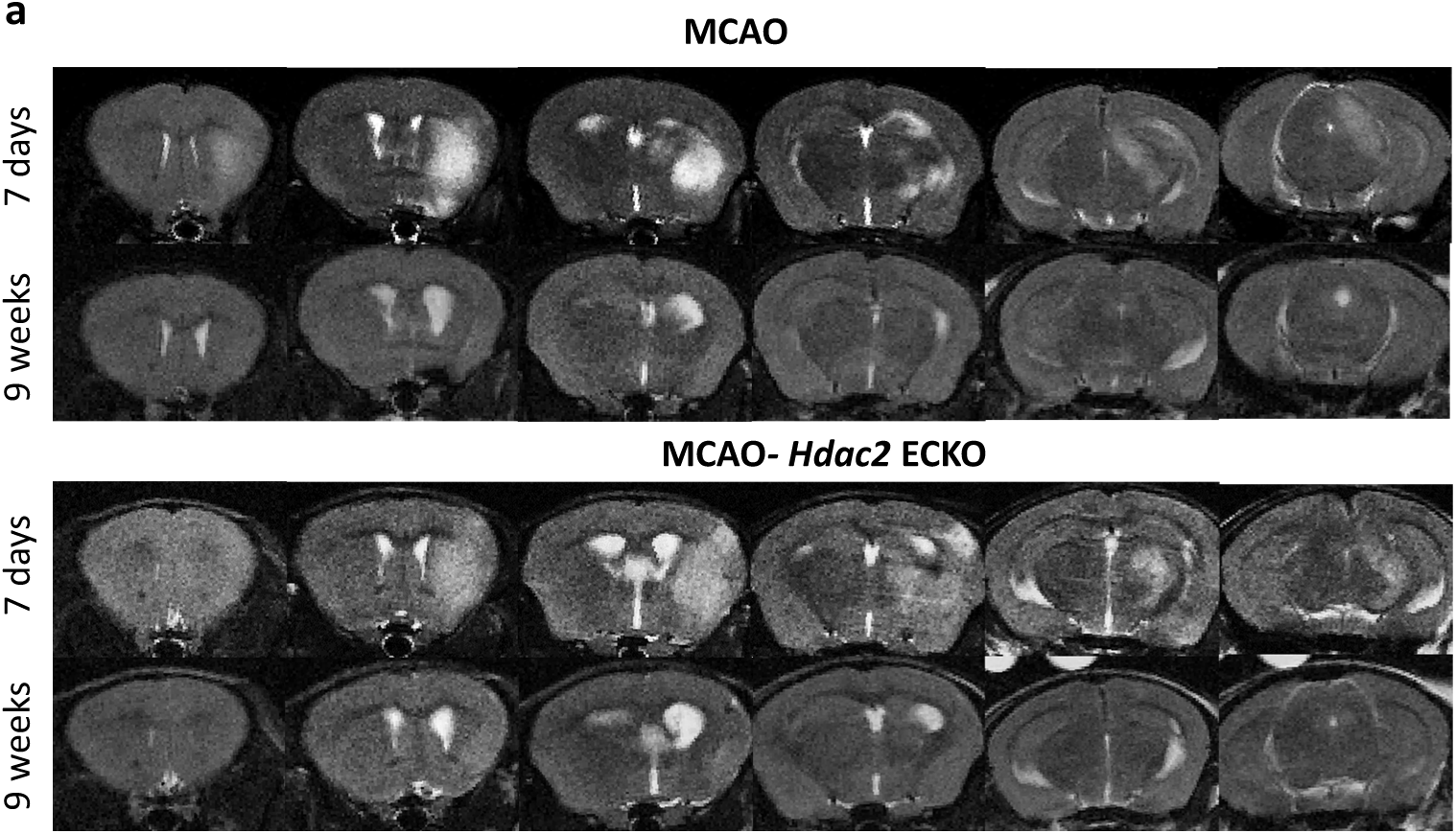
T2-weighted MRI images of MCAO and MCAO-*Hdac2* mice brains. **(a)** Six serial coronal images, arranged from rostral to caudal, depict representative MCAO and MCAO-*Hdac2* mice at 7 days and 9 weeks post-surgery. Infarct regions appear as distinct T2-hyperintense areas in the brains of both groups at 7 days post-surgery.

**Figure-8 Supplement:**
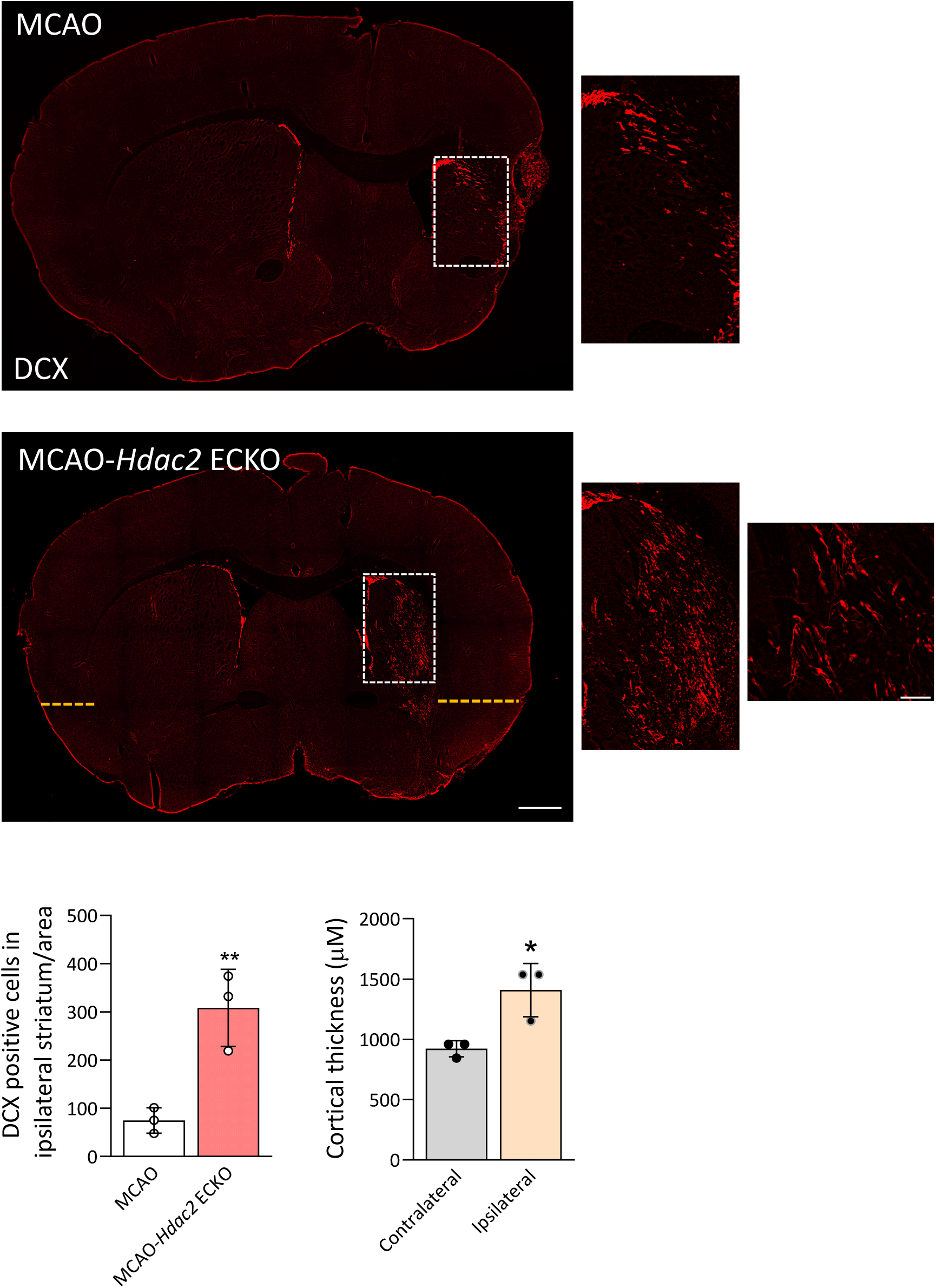
CNS EC *Hdac2* deletion augments DCX-positive immature neurone population in the stroke brain. **(a)** Representative images of DCX-stained sections in MCAO and MCAO-*Hdac2* ECKO. Areas within white dotted lines are digitally magnified to show DCX^+^ cells in MCAO and MCAO-*Hdac2* ECKO. 10x images were acquired and merged using tile scanning. Scale bar: 500 µm. 40x image from MCAO-*Hdac2* ECKO mouse displaying the pattern of distribution of DCX ^+^ cells in the ipsilateral striatum. Scale bar: 50 µm. Quantification of the number of DCX^+^ cells in the ipsilateral cortex, revealing a significant increase in MCAO-*Hdac*2 ECKO brains compared to MCAO. (**p ≤ 0.01 vs. MCAO; N =3 /group; three matching sections per mouse were analyzed). **(b)** Graph displaying quantification of cortical thickness between ipsilateral and contralateral sides of MCAO-*Hdac*2 ECKO brains. Cortical thickness was significantly increased in the ipsilateral sides of MCAO-*Hdac*2 ECKO brains. (*p ≤ 0.05 vs. MCAO; N =3 /group; matching sections per mouse were analyzed).

